# Multifaceted Hi-C benchmarking: what makes a difference in chromosome-scale genome scaffolding?

**DOI:** 10.1101/659623

**Authors:** Mitsutaka Kadota, Osamu Nishimura, Hisashi Miura, Kaori Tanaka, Ichiro Hiratani, Shigehiro Kuraku

## Abstract

**Background:** Hi-C, a derivative of chromosome conformation capture (3C) targeting the whole genome, was originally developed as a means for characterizing chromatin conformation. More recently, this method has also been frequently employed in elongating nucleotide sequences obtained by *de novo* genome sequencing and assembly, in which the number of resultant sequences rarely converge into the chromosome number. Despite the prevailing and irreplaceable use, sample preparation methods for Hi-C have not been intensively discussed, especially from the standpoint of genome scaffolding.

**Results:** To gain insights into the best practice of Hi-C scaffolding, we performed a multifaceted methodological comparison using vertebrate samples and optimized various factors during sample preparation, sequencing, and computation. As a result, we have identified some key factors that help improve Hi-C scaffolding including the choice and preparation of tissues, library preparation conditions, and restriction enzyme(s), as well as the choice of scaffolding program and its usage.

**Conclusions:** This study provides the first comparison of multiple sample preparation kits/protocols and computational programs for Hi-C scaffolding, by an academic third party. We introduce a customized protocol designated the ‘<u>i</u>nexpensive and <u>con</u>trollable <u>Hi-C</u> (iconHi-C) protocol’, in which the optimal conditions revealed by this study have been incorporated, and release the resultant chromosome-scale genome assembly of the Chinese softshell turtle *Pelodiscus sinensis*.

## Background

Chromatin, a complex of nucleic acids (DNA and RNA) and proteins, exhibits a complex three-dimensional organization in the nucleus, which enables intricate regulation of genome information expression through spatiotemporal controls (reviewed in [1]). In order to characterize chromatin conformation on a genomic scale, the Hi-C method was introduced as a derivative of chromosome conformation capture (3C) (Fig. 1A; [2]). This method detects chromatin contacts on a genomic scale through digestion of crosslinked DNA molecules with restriction enzymes, followed by proximity ligation of the digested DNA molecules. Massively parallel sequencing of the library harboring ligated DNA molecules enables comprehensive quantification of contacts between different genomic regions inside and between chromosomes, which is presented in a heatmap conventionally called the ‘contact map’ [3].

**Figure 1:**
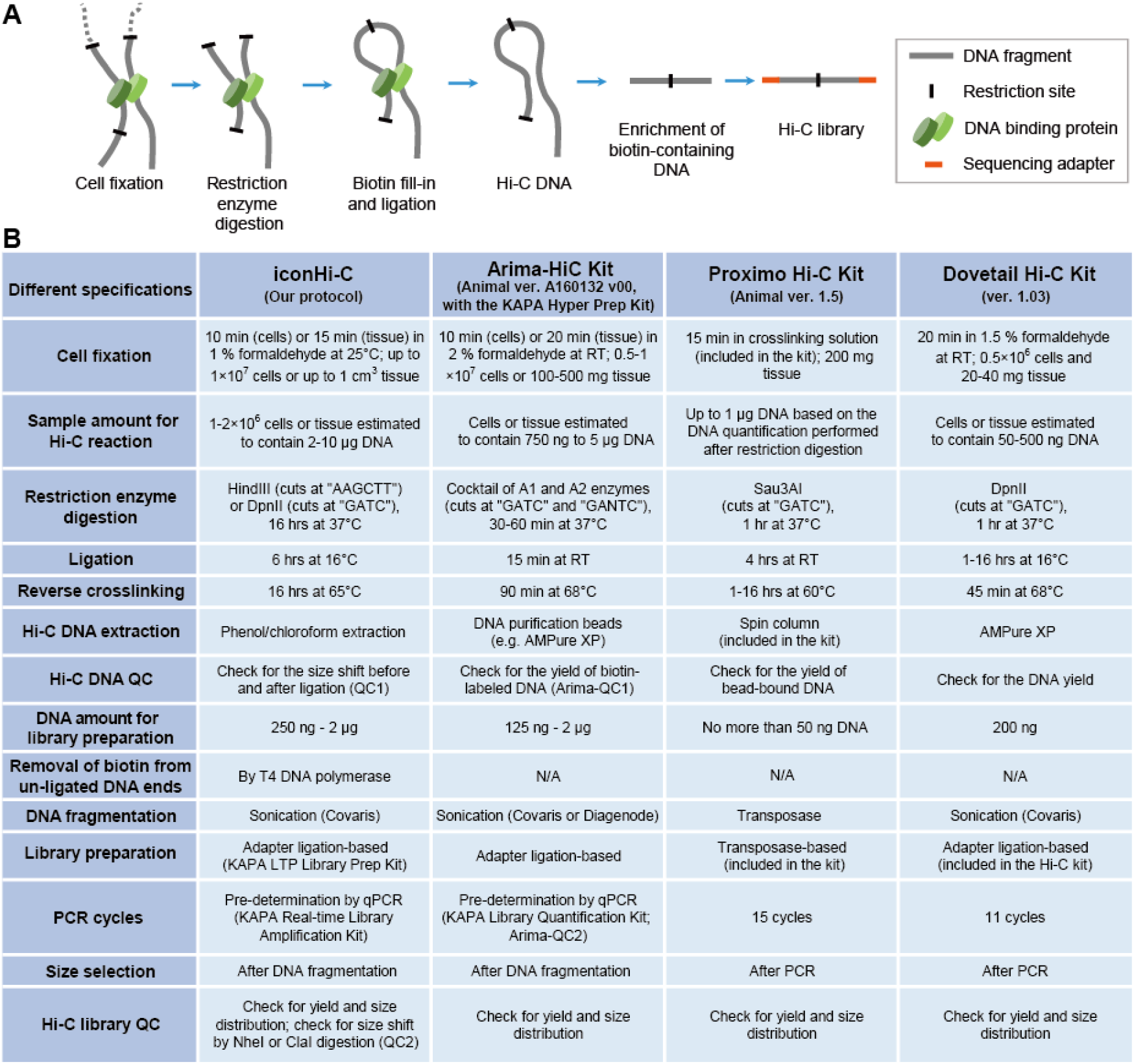
Hi-C library preparation. (A) Basic procedure. (B) Comparison of Hi-C library preparation methods. Included here are only the major differences between the methods. The KAPA Hyper Prep Kit (KAPA Biosystems) is assumed to be conjunctly used with Arima Hi-C Kit, among the several specified kits. See Supplementary Protocol S1 for the full version of the iconHi-C protocol which was derived from the protocol previously introduced [23].

Analyses of chromatin conformation with Hi-C have revealed more frequent contacts between more closely linked genomic regions, which has prompted this method to be employed in elongating *de novo* genome sequences, more recently [4]. In *de novo* genome sequencing, the number of assembled sequences is usually far larger than the number of chromosomes in the karyotype of the species of interest, irrespective of the sequencing platform chosen [5]. The application of Hi-C scaffolding enabled remarkable enhancement of sequence continuity to reach a chromosome scale and integration of fragmentary sequences into longer sequences, which are similar in number to that of chromosomes in the karyotype. In early 2018, commercial Hi-C library preparation kits were introduced to the market (Fig. 1B), and *de novo* genome assembly was revolutionized by the release of versatile computational programs for Hi-C scaffolding (Table 1), namely LACHESIS [6], HiRise [7], SALSA [8, 9], and 3d-dna [10]. These movements assisted the rise of mass sequencing projects targeting a number of species, such as Earth BioGenome Project (EBP) [11], Genome 10K (G10K)/Vertebrate Genome Project (VGP) [12, 13], and DNA Zoo Project [14]. Optimization of Hi-C sample preparation, however, has been limitedly attempted [15]. Thus, it remains unexplored which factor in particular makes a difference in the results of Hi-C scaffolding, mainly because of its costly and resource-demanding nature.

**Table 1:** Overview of the specification of the scaffolding programs released to date.

| Program | Support and availability | Input data requirement | Other information | Literature |
| --- | --- | --- | --- | --- |
| LACHESIS | Developer's support discontinued;<br>intricate installation | Generic bam format | No function to correct scaffold misjoins | [4] |
| HiRise | Open source version at GitHub<br>not updated since 2015 | Generic bam format | Employed in Dovetail Chicago/Hi-C service.<br>Default input sequence length cutoff=1000 bp | [7] |
| 3d-dna | Actively maintained and supported by the developer | Not compatible with multiple enzymes;<br>Accept only Juicer mapper format | Default parameters: -t 15000 (input sequence length cutoff), -r 2 (no. of iterations for misjoin correction) | [10, 34] |
| SALSA2 | Actively maintained and supported by the developer | Compatible with multiple enzymes;<br>generic bam (bed) file, assembly graph, unitig, 10x link files | Default parameters: -c 1000 (input sequence length cutoff), -i 3 (no. of iterations for misjoin correction) | [8, 9] |

Together with performing protocol optimization using human culture cells, we focused on the softshell turtle *Pelodiscus sinensis* (Fig. 2). This species has been adopted as a study system for evolutionary developmental biology (Evo-Devo), including the study on the formation of the dorsal shell (carapace) (reviewed in [16]). It is anticipated that relevant research communities have access to genome sequences of optimal quality. In Japan, live materials (adults and embryos) of this species are available through local farms mainly between May and August, which allows its high utility for sustainable research. Based on a previous cytogenetic report, the karyotype of this species consists of 33 chromosome pairs including Z and W (2n = 66) that show a wide variety of sizes (conventionally categorized into macrochromosomes and microchromosomes) [17]. Despite its moderate global GC-content in its whole genome at around 44%, an earlier study suggested the intragenomic heterogeneity of GC-content between and within the chromosomes, along with their sizes [18]. A wealth of cytogenetic efforts on this species accumulated fluorescence *in situ* hybridization (FISH)-based mapping data for 162 protein-coding genes covering almost all chromosomes [17–19], which serves as structural landmarks for validating genome assembly sequences.

**Figure 2:**
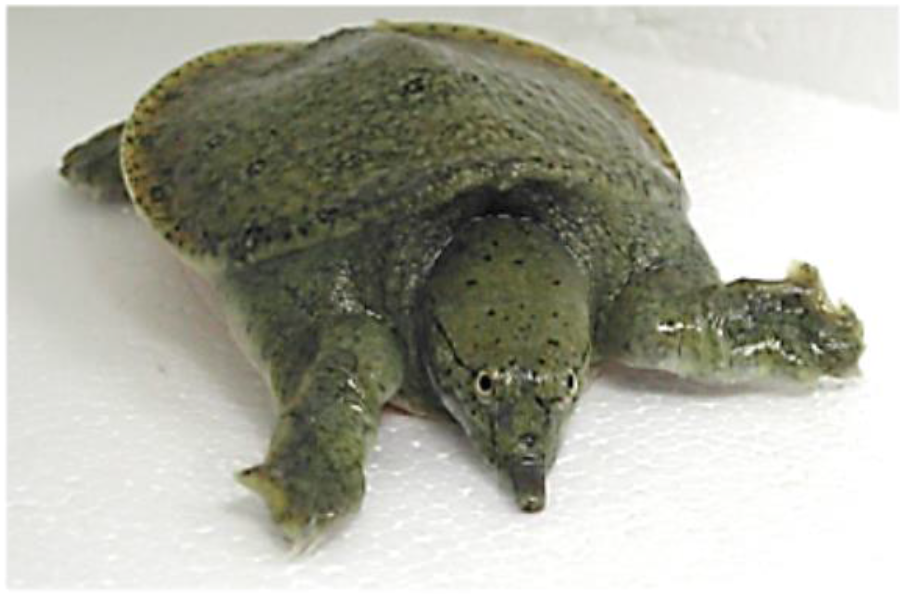
A juvenile softshell turtle *Pelodiscus sinensis*.

A draft sequence assembly of the softshell turtle genome was built with short reads and released already in 2013 [20]. This sequence assembly achieved the N50 scaffold length of >3.3 Mb but remains fragmented into approximately 20,000 sequences (see Supplementary Table S1). The longest sequence in this assembly is only slightly larger than 16 Mb, which is much shorter than the largest chromosome size estimated from the karyotype report [17]. The total size of the assembly is approximately 2.2 Gb, which is a moderate size for a vertebrate species. Because of its affordable genome size, sufficiently complex structure, and availability of validation methods, we reasoned that the genome of this species is a suitable target for our methodological comparison, and its improved genome assembly is expected to assist a wide range of genome-based studies employing this species.

## Results

### Stepwise QC before large-scale sequencing

It would be ideal to judge the quality of prepared libraries before costly sequencing. Following existing literature [15, 21], we routinely control the quality of Hi-C DNAs and Hi-C libraries by observing DNA size shifts with digestion targeting the restriction sites in properly prepared samples (Fig. 3). More concretely, a successfully ligated Hi-C DNA sample should exhibit a slight length recovery of restricted DNA fragments after ligation (QC1), which serves as an indicator of qualified samples (e.g., Sample 1 in Fig. 3B). In contrast, an unsuccessfully prepared Hi-C DNA does not exhibit this length recovery (e.g., Sample 2 in Fig. 3B). In a later step, DNA molecules in a successfully prepared HindIII-digested Hi-C library should contain the NheI restriction site at a high probability. Thus, the length distribution after the NheI digestion of the prepared library serves as an indicator of qualified or disqualified products (QC2; Fig. 3C). This series of QCs is incorporated into our protocol by default (Supplementary Protocol S1) and can also be performed along with sample preparation using commercial kits provided that it employs a single restriction enzyme.

**Figure 3:**
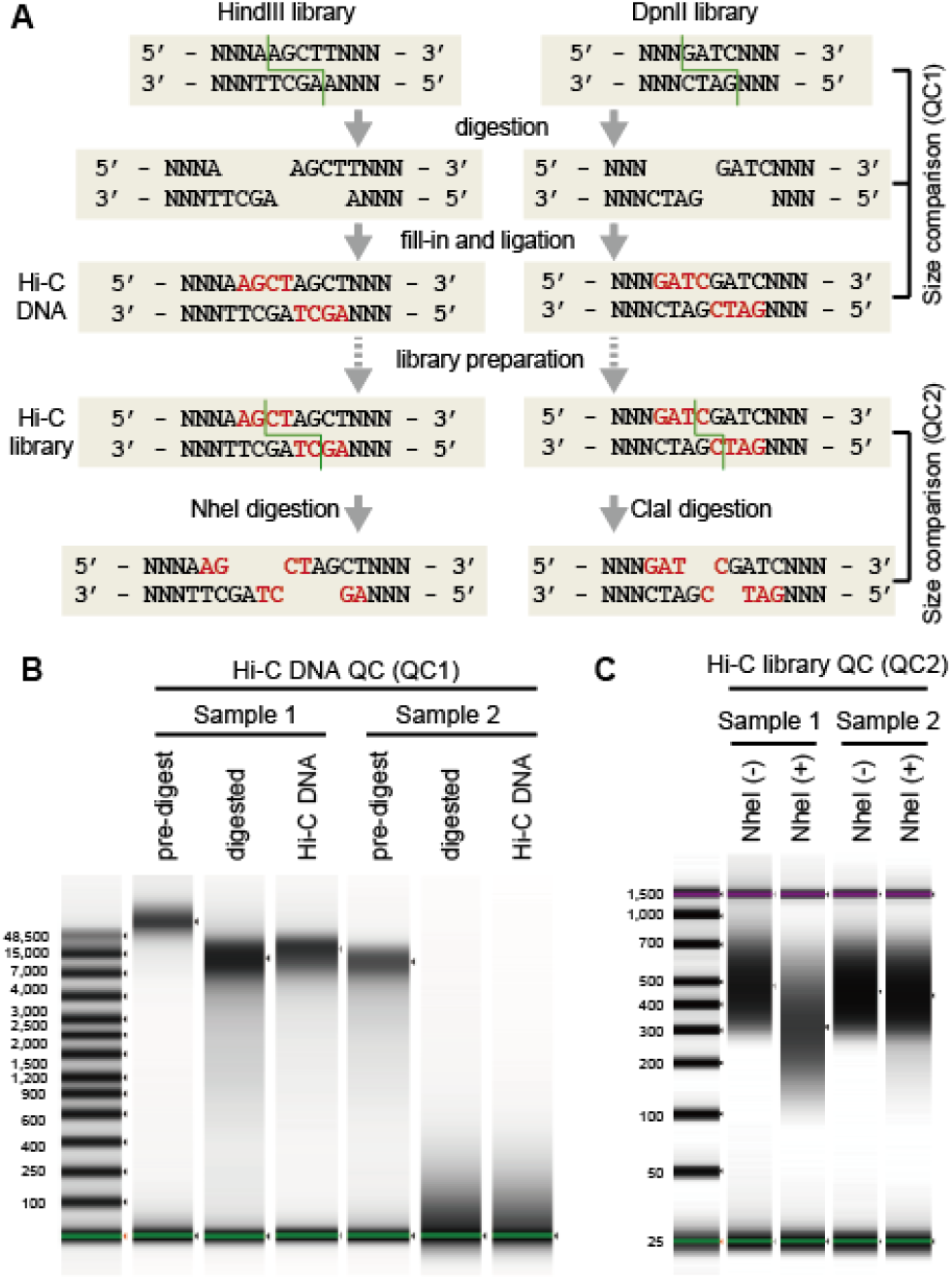
Structure of Hi-C DNA and principle of quality controls. (A) Schematic representation of the library preparation workflow based on HindIII or DpnII digestion. Patterns of restriction are indicated by the green lines. Nucleotides that were filled in are indicated by the letters in red. (B) Size shift analysis of HindIII-digested Hi-C DNA (QC1). Shown are the representative images of qualified (Sample 1) and disqualified samples (Sample 2). (C) Size shift analysis of the HindIII-digested Hi-C library (QC2). Shown are the representative images of the qualified (Sample 1) and disqualified (Sample 2) samples. Size distributions were measured with Agilent 4200 TapeStation.

Some of the libraries we have prepared passed the QC steps before sequencing but yielded an unpreferably large proportion of unusable read pairs. To identify such libraries, we routinely performed small-scale sequencing with the purpose of quick and inexpensive QC using the HiC-Pro program [22] (see Fig. 4 for the read pair categories assigned by HiC-Pro). Our test with variable input data sizes (500 K–200 M read pairs) resulted in highly similar breakdowns into different categories of read pair properties (Supplementary Table S2) and guaranteed the QC with an extremely small data size of 1 M or fewer reads. These post-sequencing QC steps that do not incur a large cost are expected to help avoid large-scale sequencing of unsuccessful libraries that have somehow passed through QC1 and QC2 steps. Importantly, libraries that have passed this QC can be further sequenced in more depth as necessary.

**Figure 4:**
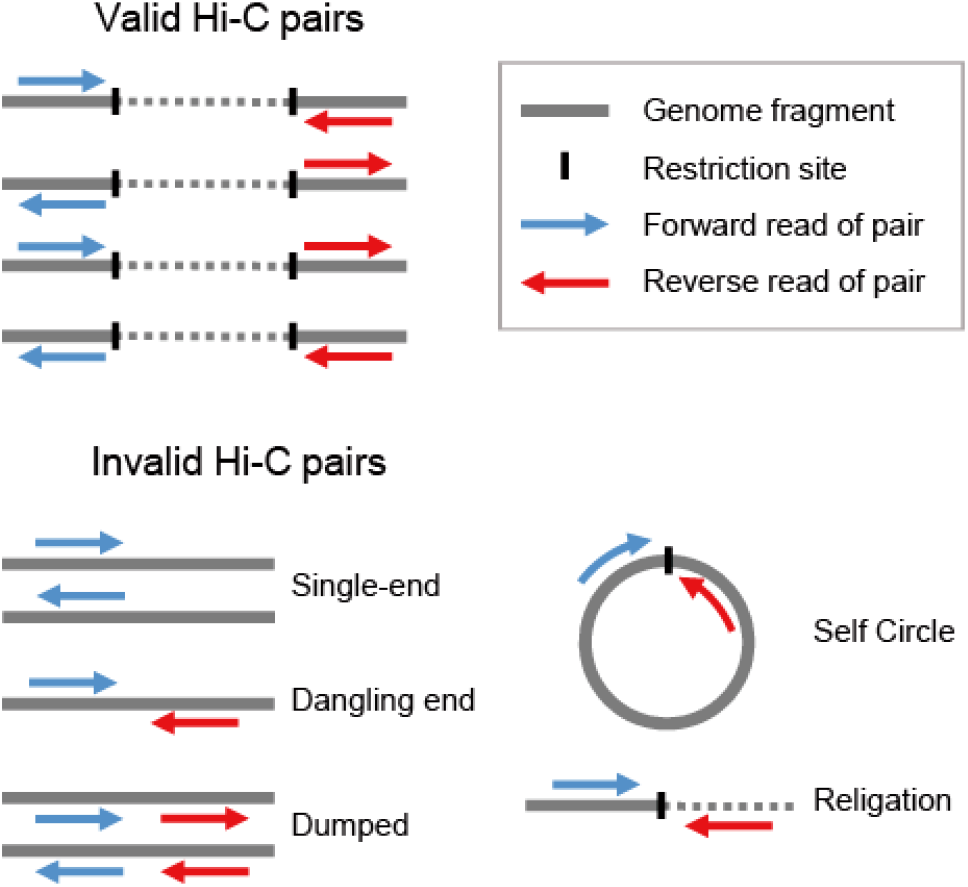
Post-sequencing quality control of Hi-C reads. Read pairs were categorized into valid and invalid pairs by HiC-Pro, based on their status in the mapping to the reference genome (see Methods). This figure was adapted from the literature originally introducing HiC-Pro [22].

## Optimization of sample preparation conditions

We identified overt differences between sample preparation protocols of already published studies and those of commercial kits (Fig. 1B). Therefore, we first sought to optimize the conditions of several preparation steps using human culture cells.

To evaluate the effect of the degree of cell fixation, we prepared Hi-C libraries from GM12878 cells fixed for 10 and 30 minutes. Our comparison did not detect any marked difference in the quality of Hi-C DNA (QC1; Fig. 5A) and Hi-C library (QC2; Fig. 5B). However, libraries with longer fixation showed larger proportions of dangling end read pairs and re-ligation read pairs, as well as a smaller proportion of valid interaction reads (Fig. 5C). Increased duration of cell fixation reduces the proportion of long-range (>1 Mb) interactions among the overall captured interactions (Fig. 5D).

**Figure 5:**
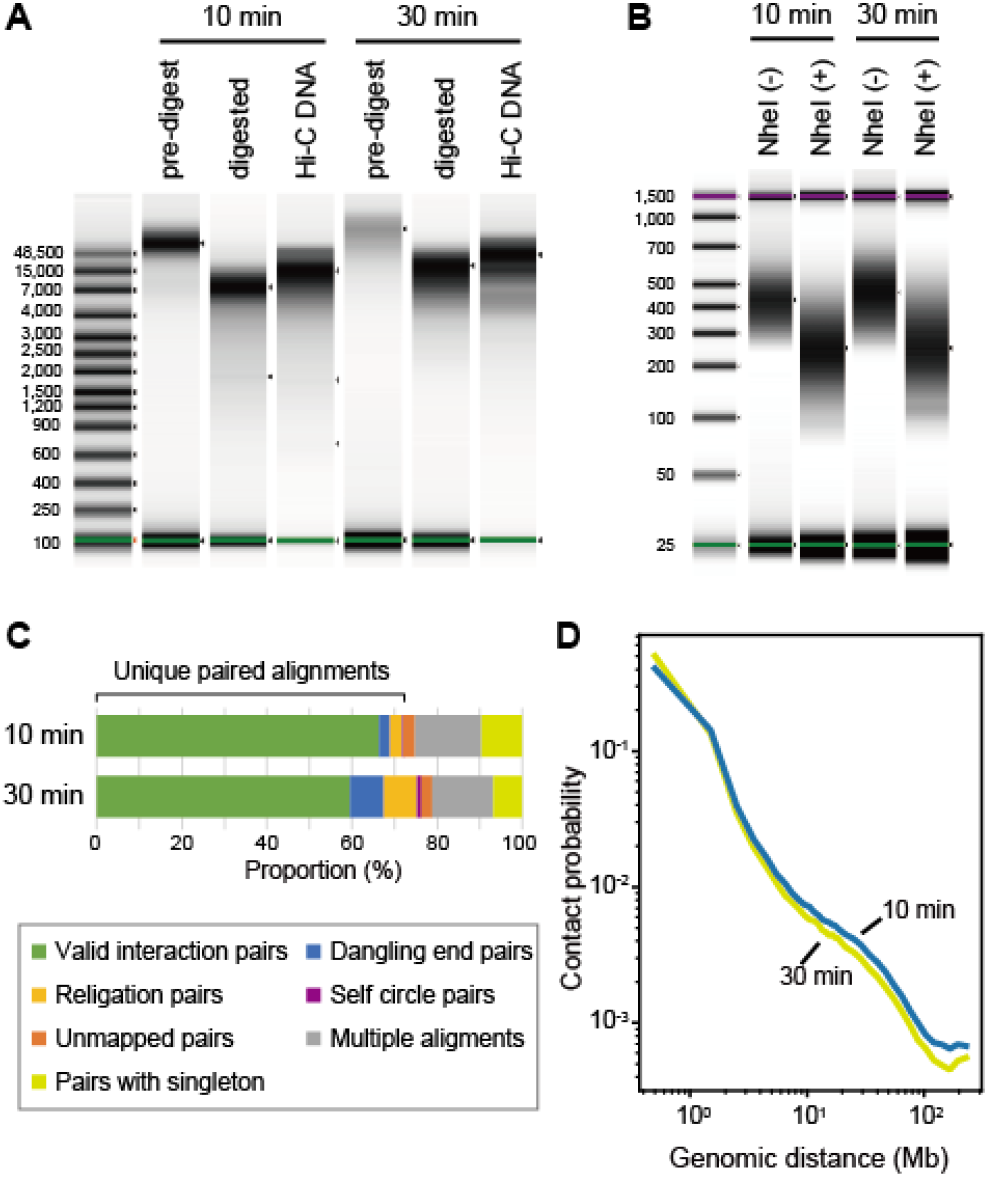
Effect of cell fixation duration. (A) QC1 of the HindIII-digested Hi-C DNA of human GM12878 cells fixed for 10 or 30 minutes in 1% formaldehyde. (B) QC2 of the HindIII-digested library of human GM12878 cells. (C) Quality control of the sequence reads by HiC-Pro using 1M read pairs. See Fig. 4 for the details of the read pair categorization. See Supplementary Table S7 for the actual proportion of the reads in each category. (D) Contact probability measured by the ratio of observed and expected frequencies of Hi-C read pairs mapped along the same chromosome [43].

The reduced preparation time with commercial Hi-C kits (up to two days according to their advertisement) is attributable mainly to shortened duration of restriction and ligation (Fig. 1B). To monitor the effect of shortening these enzymatic reactions, we analyzed the progression of restriction and ligation in a time course experiment using human GM12878 cells. The results show persistent progression of restriction until 16 hours and of ligation until 6 hours (Fig. 6).

**Figure 6:**
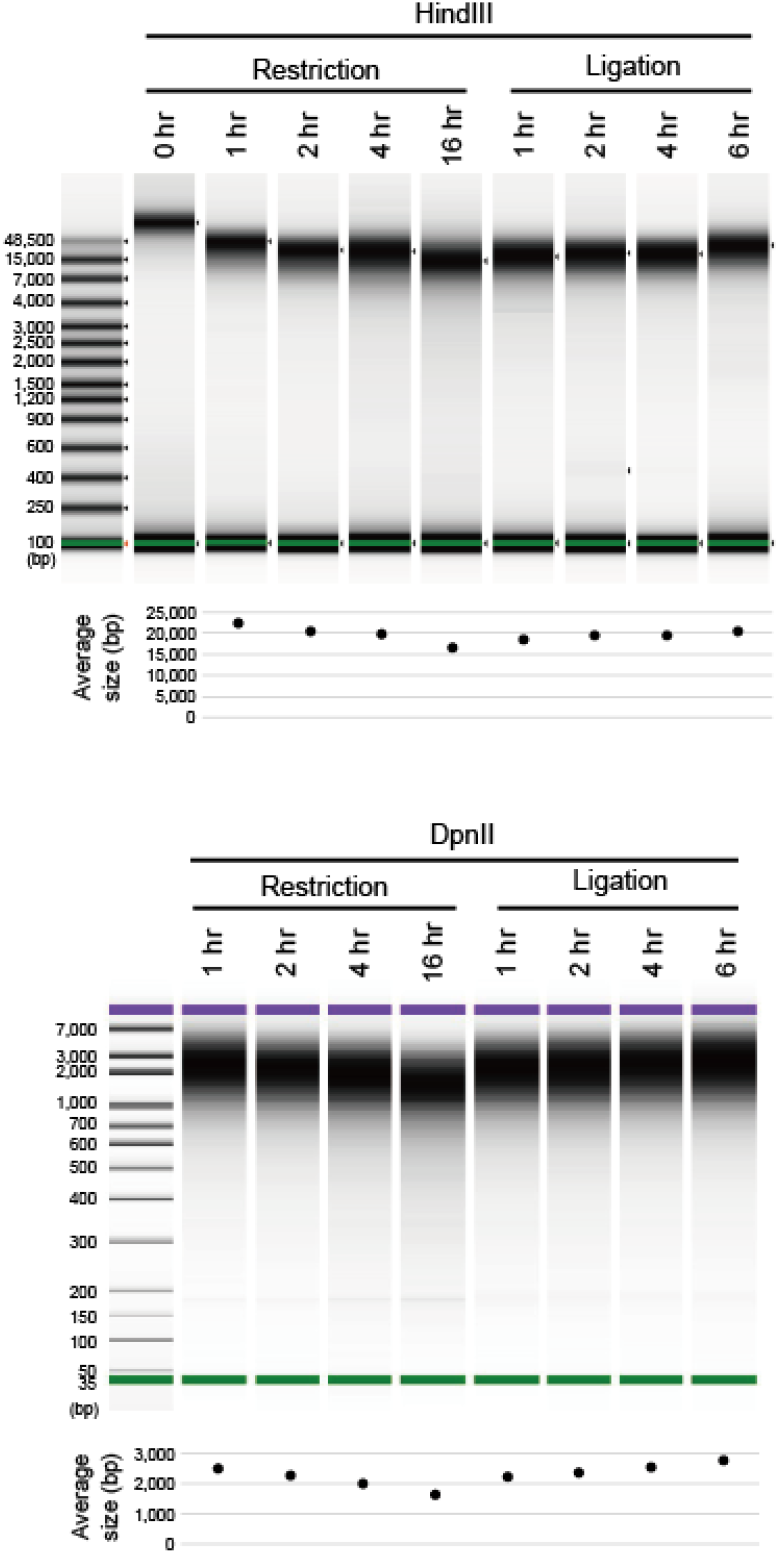
Testing variable durations of restriction and ligation of Hi-C DNA. Length distributions of the DNA molecules prepared from human GM12878 cells after variable durations of restriction and ligation are shown. Size distribution for the HindIII-digested samples (top) and DpnII-digested samples (bottom) were measured by Agilent 4200 TapeStation and Agilent Bioanalyzer, respectively.

### Multifaceted comparison using softshell turtle samples

On the basis of the detailed optimization of sample preparation conditions described above, we built an original protocol, designated the ‘iconHi-C protocol’, with 10 min-long cell fixation, 16 hour-long restriction, 6 hour-long ligation, and successive QC steps (Methods; also see Supplementary Protocol S1; Fig. 1B).

We performed Hi-C sample preparation and scaffolding using tissues from a female Chinese softshell turtle which is known to have both Z and W chromosomes [17]. For this purpose, we prepared Hi-C libraries with variable tissues (liver or blood cells), restriction enzymes (HindIII or DpnII), and protocols (our iconHi-C protocol, the Arima Genomics kit in conjunction with the KAPA Hyper Prep Kit, or the Phase Genomics kit) as outlined in Fig. 7A (see Supplementary Table S3; Supplementary Fig. S1). As in some existing protocols (e.g., [23]), we performed T4 DNA polymerase treatment in our iconHi-C protocol (Library a–d), expecting reduced proportions of ‘dangling end’ read pairs that contain no ligated junction and thus do not contribute to Hi-C scaffolding. We also incorporated this T4 DNA polymerase treatment in the workflow of the Arima kit (Library e vs. Library f without this additional treatment). We also tested a lesser degree of PCR amplification (11 cycles) along with the use of the Phase Genomics kit which compels as many as 15 cycles by default (Library h vs. Library g; Fig. 7A).

**Figure 7:**
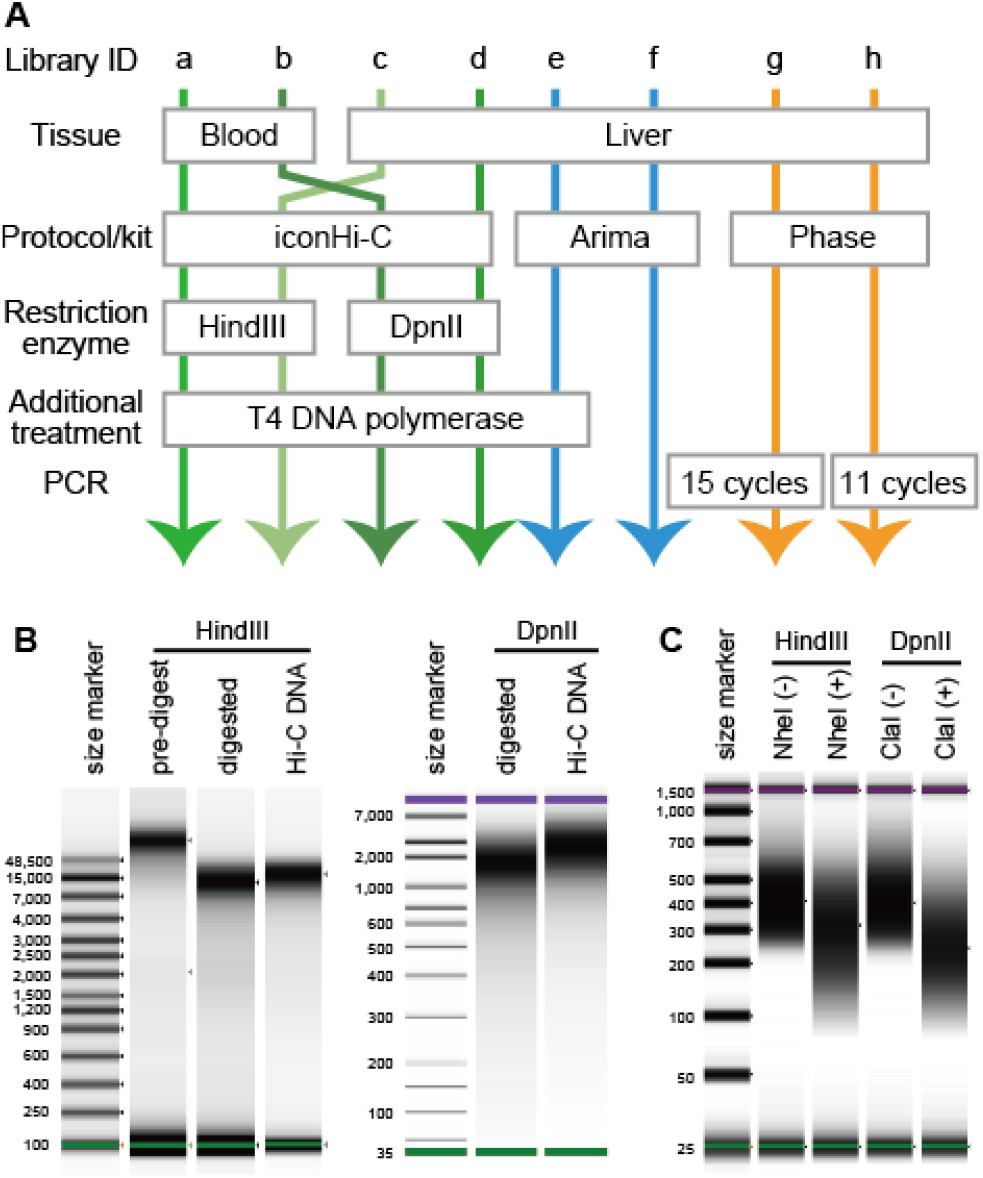
Softshell turtle Hi-C libraries prepared for our methodological comparison. (A) Lineup of the prepared libraries. This chart includes only the conditions that varied preparation methods between these libraries, and the rest of the preparation workflows are described in Supplementary Protocol S1 for the non-commercial (‘iconHi-C’) protocol and the manuals of the commercial kits. (B) Quality control of Hi-C DNA (QC1) for Library c and d. The prepared Hi-C DNA for the Chinese softshell turtle liver samples were digested with either HindIII or DpnII. (C) Quality control of Hi-C libraries (QC2). The prepared softshell turtle liver HindIII library was digested by NheI, and the DpnII library was digested by ClaI (see Fig. 3 for the technical principle). See Supplementary Fig. S3 for the QC1 and QC2 results for the samples prepared from the blood of this species.

The samples prepared with the iconHi-C protocol, which is compatible with the abovementioned QC1 and QC2, were all judged as qualified, by these QCs (Fig. 7B). The prepared Hi-C libraries were sequenced to obtain one million 127nt-long read pairs and subjected to post-sequencing QC with the HiC-Pro program (Fig. 8). As a result of this QC, the largest proportion of ‘valid interaction’ pairs was observed for Arima libraries (Library e and f). As for the iconHi-C libraries (Library a–d), fewer ‘unmapped’ and ‘religation’ pairs were detected with the DpnII libraries than with HindIII libraries. It should be noted that the QC results for the softshell turtle libraries generally produced lower proportions of the ‘valid interaction’ category and larger proportions of ‘unmapped pairs’ and ‘pairs with singleton’ than those for human libraries. This cross-species difference is accounted for by possibly incomplete genome sequences used as a reference for Hi-C read mapping (Supplementary Table S1). This evokes a caution in comparing QC results across species.

**Figure 8:**
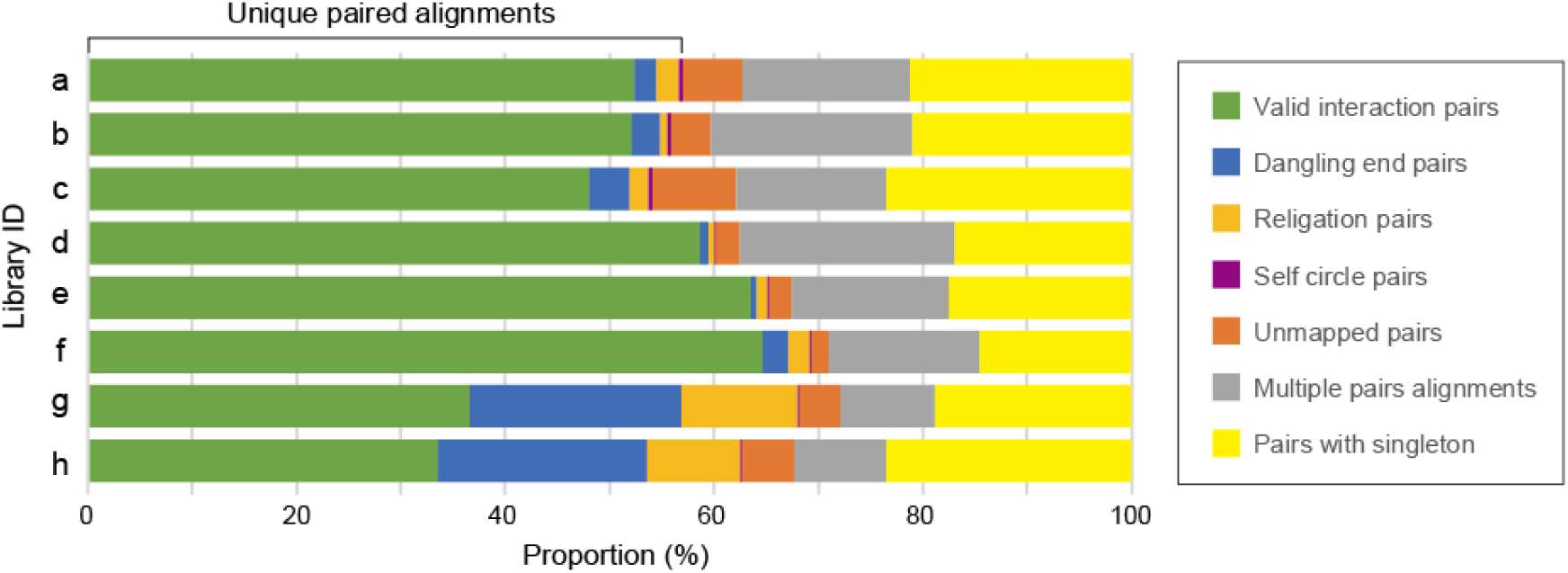
Results of the post-sequencing quality control with HiC-Pro. One million read pairs were used for computation with HiC-Pro. See Fig. 7A for the preparation conditions of Library a-h, Fig. 4 for the categorization, and Supplementary Table S3 for the actual proportion of the reads in each category. Post-sequencing quality control using variable read amounts (500 K–200 M pairs) for one of these softshell turtle libraries (Supplementary Table S6) and human GM12878 libraries (Supplementary Table S2) shows the validity of this quality control with as few as 500 K read pairs.

### Scaffolding with variable inputs and computational conditions

In this study, only well-maintained, open-source programs, namely 3d-dna and SALSA2, were used in conjunction with variable combinations of an input library, an input read amount, an input sequence cutoff length, and a number of iterative misjoin correction rounds (Fig. 9A). As a result of scaffolding, we observed a wide spectrum of basic metrics, including the N50 scaffold length (0.6–303 Mb), the largest scaffold length (8.7–703 Mb), and the number of chromosome-sized (>10 Mb) sequences (0–65) (Fig. 9; Supplementary Table S4).

**Figure 9:**
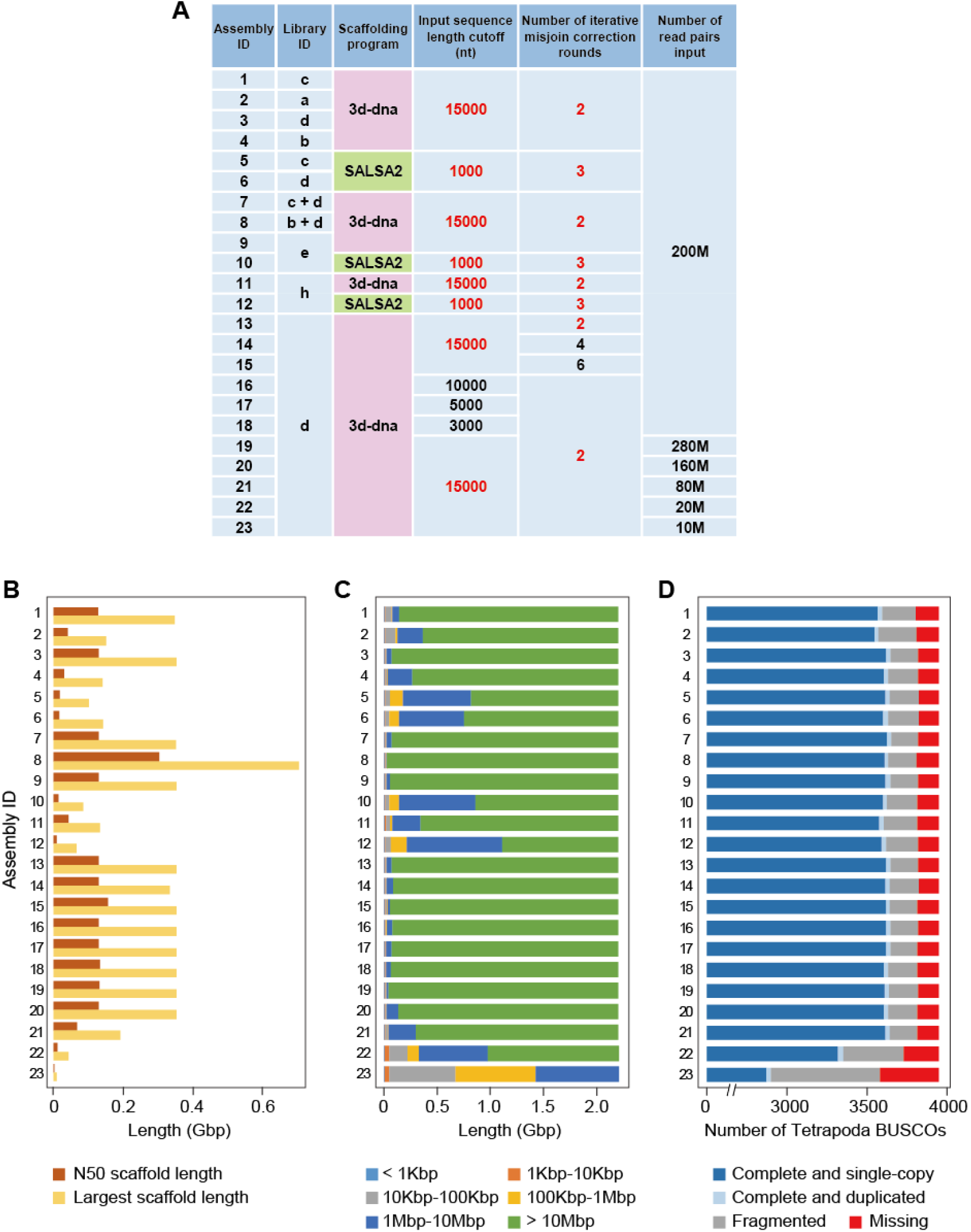
Comparison of Hi-C scaffolding products. (A) Scaffolding conditions to produce Assembly 1 to 23. Default parameters are shown with red letters. (B) Total and N50 scaffold lengths. (C) Scaffold length distributions. (D) Gene space completeness. See the panel A for Library IDs and Supplementary Table S4 for raw values of the metrics in B–D.

First of all, with the default parameters, 3d-dna consistently produced more continuous assemblies than SALSA2 (see Assembly 1 vs. 5, 3 vs. 6, 9 vs. 10, and 11 vs 12 in Fig. 9). Second, increasing the number of iterative corrections (‘-r’ option with 3d-dna) resulted in relatively large N50 lengths but with more missing orthologs (see Assembly 13–15). Third, a smaller input sequence cutoff length (‘-i’ option with 3d-dna) resulted in a smaller number of resultant scaffolds but again, with more missing orthologs (see Assembly 13, 16–18). Fourth, using the liver libraries consistently resulted in a higher continuity than using the blood cell libraries (see Assembly 1 vs. 2 as well as 3 vs. 4 in Fig. 9).

Of those, Assembly 8, employing input Hi-C reads derived from both liver and blood, exhibited an outstandingly large N50 scaffold length (303 Mb) but a larger number of undetected reference ortholog (141 orthologs) than most of the other assemblies. The largest scaffold (scaffold 5) in this assembly is approximately 703 Mb long, causing the large N50 length, and accounts for approximately one-third of the whole genome in length, as a result of possible overassembly bridging 14 putative chromosomes (see Supplementary Fig. S2).

The choice of restriction enzymes has not yet been discussed in depth, in the context of genome scaffolding. In the present study, we separately prepared Hi-C libraries with HindIII and DpnII. We did not mix multiple enzymes in a reaction (apart from using the Arima kit originally employing two enzymes) and instead performed a single scaffolding run with both HindIII-based and DpnII-based reads (see Assembly 7 in Fig. 9). Our comparison of multiple metrics expectedly highlights a more successful result with DpnII than with HindIII (see Assembly 1 vs. 3 as well as 2 vs. 4; Fig. 9). However, the mixed input of HindIII-based and DpnII-based reads did not necessarily yield a better scaffolding result (see Assembly 3 vs. 7).

### Validation of scaffolding results with transcriptome and FISH data

In addition to the above-mentioned evaluation of the scaffolding results based on sequence length and gene space completeness, we attempted to evaluate the sequence continuity with independently obtained data. First, we mapped assembled transcript sequences onto our Hi-C scaffold sequences (see Methods). This did not reveal any substantial differences between the assemblies (Supplementary Table S5), probably because the sequence continuity after Hi-C scaffolding already exceeded that of RNA-seq library inserts even when the lengths of intervening introns in the genome are taken into consideration. The present analysis with RNA-seq data did not provide an effective resort of continuity validation.

Second, we referred to the fluorescence *in situ* hybridization (FISH) mapping data for 162 protein-coding genes from published cytogenetic studies [17–19], which allowed us to check the locations of those genes with our resultant Hi-C assemblies. In this analysis, we evaluated Assembly 3, 7, and 9 (see Fig. 9A) that showed better scaffolding results in terms of sequence length distribution and gene space completeness (Fig. 9B). As a result, we confirmed the positioning of almost all genes and their continuity over the centromeres, which encompassed not only large but also small chromosomes (conventionally called ‘macro-’ and ‘micro-chromosomes’; Fig. 10). Two genes that were not confirmed by Assembly 7 (*UCHL1* and *COX15*; Fig. 10) were found in separate scaffold sequences shorter than 1 Mb, which indicates insufficient scaffolding. On the other hand, the gene array including *RBM5*, *TKT*, *WNT7A*, and *WNT5A*, previously shown by FISH, was consistently unconfirmed by all the three assemblies (Fig. 10), which did not provide any clue for among-assembly evaluation or even indicated an erroneous interpretation of FISH data in a previous study.

**Figure 10:**
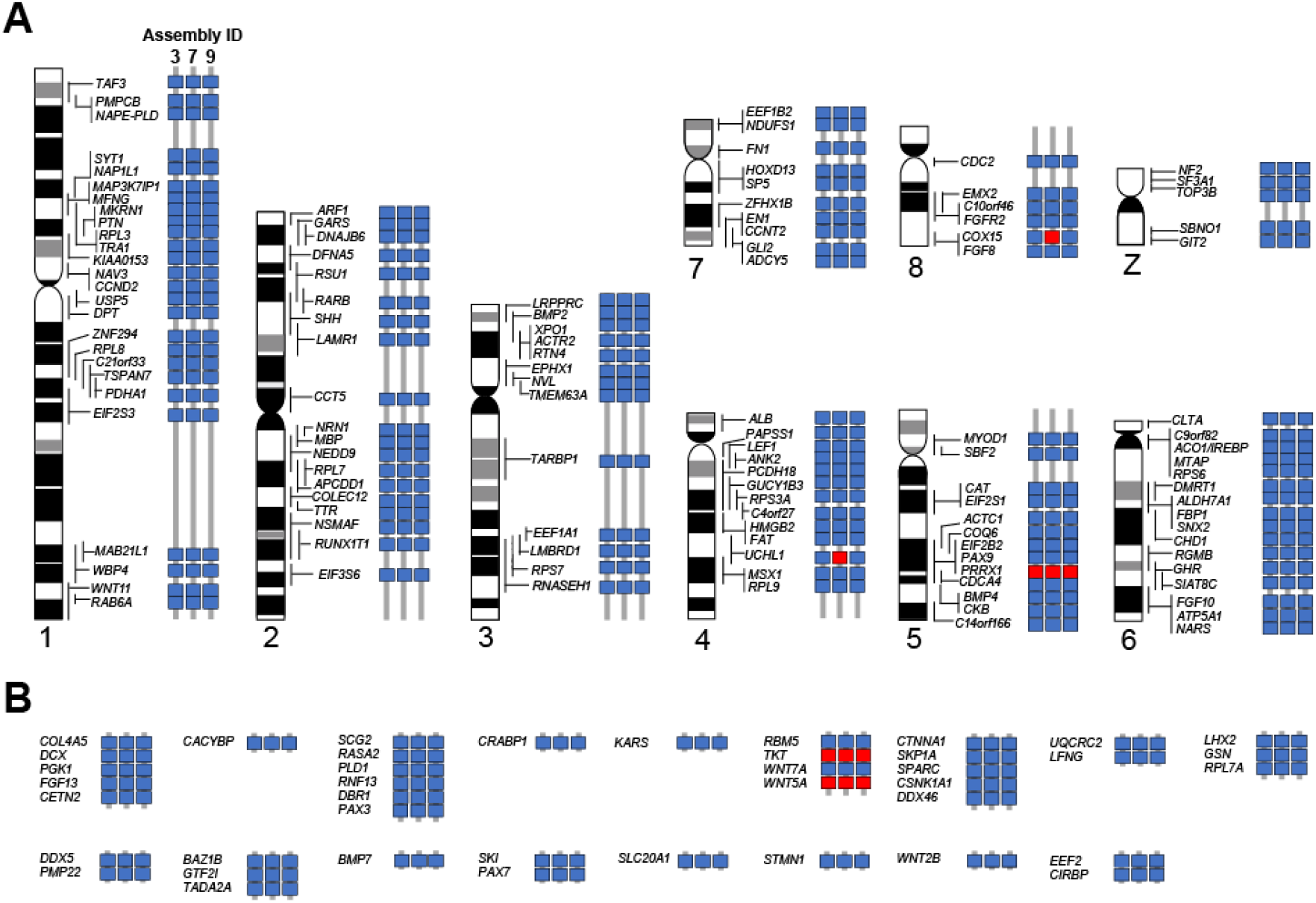
Cytogenetic validation of Hi-C scaffolding results. On the scaffolded sequences of Assembly 3, 7, and 9, we evaluated the consistency of the positions of the selected genes that were previously localized on 8 macrochromosomes and Z chromosome (A) and microchromosomes (B) by chromosome FISH [17–19] (see Results). Concordant and discordant gene locations on individual assemblies are indicated with blue and red boxes, respectively. The arrays of genes without idiograms in B were identified on chromosomes that are cytogenetically indistinguishable from each other.

## Discussion

### Starting materials: not genomic DNA extraction but *in situ* cell fixation

In genome sequencing, best practices for high molecular weight DNA extraction have often been discussed (e.g., [24]). This factor is fundamental to building longer contigs, whether employing short-read or long-read sequencing platforms. Also, the proximity ligation method using Chicago libraries provided by Dovetail Genomics which is based on *in vitro* chromatin reconstruction [7], uses genomic DNA as starting materials. Instead, proximity guided assembly enabled by Hi-C employs cellular nuclei preserving chromatin conformation, which brings a new technical challenge for appropriate sampling and sample preservation in genomics.

In preparing the starting materials, it seems important to optimize the degree of cell fixation depending on your sample choice, to obtain an optimal result in Hi-C scaffolding (Fig. 5). Another practical lesson about tissue choice was obtained by examining Assembly 8 (Fig. 9A). This assembly was produced by 3d-dna scaffolding with both liver and blood libraries (Library b and d), which led to an unacceptable result possibly caused by overassembly (Fig. 9B–D; also see Results). It is likely that enhanced cellular heterogeneity, possibly introducing excessive conflicting chromatin contacts, did not allow the scaffolding program to properly group and order the input genome sequences. In brief, we recommend the use of samples with modest cell-type heterogeneity amenable to thorough fixation.

### Considerations in sample preparation

In this study, we could not test all commercial Hi-C kits available in the market. This is partly because the Dovetail Hi-C kit specifies a non-open source program HiRise as the only supported downstream computation solution and does not allow a direct comparison with other kits, namely those from Phase Genomics and Arima Genomics.

According to our calculation, it would be at least three times more economical to prepare a Hi-C library with the iconHi-C protocol than with a commercial kit. Practically, the cost difference would be even larger, either when one cannot fully consume the purchased kit or when one cannot undertake post-sequencing computation steps and thus cover additional outsourcing cost for this.

Genomic regions targeted by Hi-C are determined by the choice of restriction enzymes. Theoretically, 4-base cutters (e.g., DpnII), potentially with more frequent restriction sites on the genome, are expected to provide a higher resolution than 6-base cutters (e.g., HindIII) [15]. However, it might not be so straightforward when the species-by-species variation of GC-content, as well as its intra-genomic heterogeneity, are taken into consideration. The use of multiple enzymes in a single reaction could be promising, but not all scaffolding programs are compatible with multiple enzymes from a computational viewpoint (see Table 1 for a comparison of scaffolding program specifications). Another technical downside is the incompatibility of DNA ends restricted by multiple enzymes, with restriction-based QCs, such as QC2 in our iconHi-C protocol (Fig. 3). Therefore, in this study, DpnII and HindIII were separately employed in conjunction with the iconHi-C protocol, which resulted in higher scaffolding performance with the DpnII library (Figs. 8 and 9), as expected. In addition, we input the separately prepared DpnII and HindIII libraries together in scaffolding (Assembly 7), but this attempt did not lead to higher scaffolding performance (Figs. 9B–D and 10). The Arima Hi-C kit employs two different enzymes that can produce much more combinations of restriction sites, because one of the two enzymes recognizes the nucleotide stretch GANTC. Scaffolding with the libraries prepared using this kit resulted in one of the most acceptable assemblies (Assembly 9). However, this result did not explicitly exceed the performance of scaffolding with the iconHi-C libraries including the one employing only a single enzyme DpnII (Library d).

One concern about the use of commercial kits (except the Arima Hi-C kit used with the Arima-QC2) is overamplification by PCR, as their manuals specify certain numbers of PCR cycles *a priori* (15 cycles for the Phase Genomics Proximo Hi-C kit and 11 cycles for the Dovetail Hi-C kit). In our iconHi-C protocol, an optimal number of PCR cycles is estimated by means of a preliminary real-time PCR using a small aliquot (Step11.25–29 in Supplementary Protocol S1) as traditionally performed for other library types (e.g., [25]). This procedure allowed us to minimize the PCR cycles down to five cycles (Supplementary Table S3). The Dovetail Hi-C kit recommends that one consumes larger amounts of kit components than specified for a single sample, depending on the genome size, as well as the degree of genomic heterozygosity and repetitiveness, of the species of interest. However, with our iconHi-C protocol, we always performed a single library preparation, irrespective of those species-specific factors, which we understand suffices in all the cases we have tested.

Commercial Hi-C kits, usually advertised for easiness and quickness, have largely shortened the protocol down to two days, in comparison with existing non-commercial protocols (e.g., [15, 23]). Such time-saving protocols are achieved mainly by shortened durations of restriction enzyme digestion and ligation (Fig. 1B). Our assessment, however, showed unsaturated reaction within such shortened time frames employed in the commercial kits (Fig. 6). Also, our attempt to insert a step for T4 DNA polymerase treatment in sample preparation with the Arima Hi-C kit resulted in reduced ‘dangling end’ reads (Library e vs. Library f in Fig. 8). As for the Phase Genomics Proximo Hi-C kit, transposase-based library preparation contributes largely to shortening its protocol, but this decreases the operability of library insert lengths. Especially if Hi-C sample preparation is performed for a limited number of samples, as practiced typically for genome scaffolding, one would opt to consider these points, even in using commercial kits, in order to further improve the quality of prepared libraries and scaffolding products.

### Considerations in sequencing

The quantity of Hi-C read pairs to be input for scaffolding is critical because it accounts for the majority of the cost of Hi-C scaffolding. Our protocol introduces a thorough safety system to prevent sequencing unsuccessful libraries, firstly with pre-sequencing QCs for size shift analysis (Fig. 3) and secondly with small-scale (down to 500 K read pairs) sequencing (see Results; also see Supplementary Table S2, S6).

Our comparison shows a dramatic decrease in assembly quality when less than 100 M read pairs were used (see the comparison among Assembly 19–23 above in Fig. 9). Still, we obtained optimal results with a smaller number of reads (ca. 160 M per 2.2 Gb genome) than recommended by commercial kits (e.g., 100 M per 1 Gb genome for the Dovetail Hi-C kit and 200 M per Gb genome for the Arima Hi-C kit). As generally and repeatedly discussed, the proportion of informative reads and their diversity, rather than just the number of all obtained reads, are critical.

In terms of read length, we did not perform any comparison in this study. Longer reads may enhance the fidelity in characterizing the read pair property and allows precise QC. Still, the existing Illumina sequencing platform has enabled economical acquisition of 150 nt-long paired-end reads, which did not prompt us to vary the read length.

### Considerations in computation

In this study, 3d-dna produced a more reliable scaffolding output than SALSA2, whether sample preparation employed a single or multiple enzyme(s) (Fig. 9B–D). On the other hand, 3d-dna needed more time to complete scaffolding than SALSA2. Apart from the choice of the program, there are quite a few points to consider, in order to achieve successful scaffolding for a smaller investment. In general, it is advised not to take Hi-C scaffolding results for granted, and it is necessary to improve them by referring to contact maps, using an interactive tool such as Juicebox [14]. In this study, however, we compared raw scaffolding outputs to evaluate sample preparation and reproducible computational steps.

Our study employed variable parameters of the scaffolding programs (Fig. 9A). First, available Hi-C scaffolding programs have different default length cut-off values for input sequences (e.g., 15000 bp for the parameter ‘-i’ with 3d-dna and 1000 bp for the parameter ‘-c’ with SALSA2). Only sequences longer than the cut-off length value contribute to sequence elongation towards the chromosome sizes, and those shorter than that are implicitly excluded from the scaffolding process and remain unchanged. Typically with the Illumina sequencing platform, genomic regions with unusually high frequencies of GC-content and repetitive elements are not assembled into sequences with sufficient lengths (see [26]). Such genomic regions tend to be excluded from chromosome-scale Hi-C scaffolds because their length is smaller than the threshold. It is also possible that such regions are excluded because few Hi-C read pairs are mapped to such regions, even if they exceed the cutoff length. One needs to deliberately set the length cutoff in accordance with the overall continuity of the input assembly and possible interest into particular, fragmentary sequences expected to be elongated. It should be warned that lowering the length threshold can result in frequent misjoins in the scaffolding output (Fig. 9B–D) or too much computational time. Regarding the number of iterative misjoin correction rounds (the parameter ‘-r’ with 3d-dna and ‘i’ with SALSA2), our attempts with increased values did not necessarily yield favorable results (Fig. 9B–D), which did not provide a consistent optimal range of values but rather suggests the importance of performing multiple scaffolding runs with varied parameters.

### Considerations in assessing chromosome-scale genome sequences

Our assessment with cytogenetic data confirmed the continuity of gene linkage over the obtained chromosome-scale sequences (Fig. 10). This validation was necessitated by almost saturated scores of typical gene space completeness assessment such as BUSCO (Supplementary Table S4) as well as transcript contig mapping (Supplementary Table S5), both of which did not provide an effective metric for evaluation.

For further evaluation of our scaffolding results, we referred to sequence length distribution of the genome assemblies of other turtle species that are regarded as chromosome-scale. This showed comparable values for the basic metrics to our Hi-C scaffolding results on the softshell turtle, that is, a N50 length of 127.5 Mb and the maximum sequence length of 344.5 Mb for the green sea turtle (*Chelonia mydas*) genome assembly released by the DNA Zoo Project and a N50 length of 131.6 Mb and the maximum length of 370.3 Mb for the Goode’s thornscrub tortoise (*Gopherus evgoodei*) genome assembly released by the Vertebrate Genome Project (VGP). Scaffolding results should be evaluated by referring to an estimate N50 length and the maximum length based on the actual number and the length distribution of chromosomes in the intrinsic karyotype of the species in question or its close relative. Turtles tend to have the N50 length of approximately 130 Mb and the maximum length of 350 Mb, while many teleost fish genomes exhibit an N50 length of as low as 20–30 Mb and the maximum length of <100 Mb [27]. If these metrics show excessive values, scaffolded sequences harbor overassembly that erroneously boosts length-based metrics. Larger values that researchers conventionally regard as signs for successful sequence assembly do not necessarily indicate higher precision.

The total length of assembly sequences is expected to increase after Hi-C scaffolding, because scaffolding programs simply insert a stretch of the unassigned base ‘N’ with a uniform length between input sequences in most cases (500 bp as default with both 3d-dna and SALSA2). However, this has a minor impact on the total assembly sequence length. In fact, inserting the ‘N’ stretches of arbitrary lengths has been an implicit, rampant practice even before Hi-C scaffolding prevailed―for example, the most and second most frequent lengths of the ‘N’ stretch in the publicly available zebrafish genome assembly Zv10 are 100 and 10 bp, respectively.

### Conclusions

In this study, we introduced the iconHi-C protocol in which successive QC steps are implemented, and assessed possible keys for improving Hi-C scaffolding. Overall, our study shows that a small variation in sample preparation or computation for scaffolding can have a large impact on scaffolding output, and any scaffolding output should ideally be validated by independent information, such as cytogenetic data, long reads, or genetic linkage maps. Our present study aimed to evaluate the output of reproducible computational steps, which in practice should be followed by modifying the raw scaffolding output by referring to independent information or by analyzing chromatin contact maps. The study employed only limited combinations of species, sample prep methods, scaffolding programs, and its parameters, and we will continue testing different conditions for kits/programs that did not necessarily perform well here with our specific materials.

## Methods

### Initial genome assembly sequences

The softshell turtle (*Pelodiscus sinensis*) assembly published previously [20] was downloaded from NCBI GenBank (GCA_000230535.1), whose gene space completeness and length statistics were assessed by gVolante [28] (see Supplementary Table S1 for the assessment results). Although it could be suggested to remove haplotigs before Hi-C scaffolding [29], we omitted this step because of the low frequency of the reference orthologs with multiple copies (0.72 %; Supplementary Table S1), indicating a minimal degree of haplotig contamination.

## Animals and cells

We sampled tissues (liver and blood cells) from a female purchased from a local farmer in Japan, because the previous whole genome sequencing used the whole blood of a female [20]. All the experiments were conducted in accordance with the Guideline of the Institutional Animal Care and Use Committee of RIKEN Kobe Branch (Approval ID: A2017-12).

Human lymphoblastoid cell line GM12878 was purchased from the Coriell Cell Repositories and cultured in RPMI-1640 media (Thermo Fisher Scientific) supplemented with 15% FBS, 2 mM L-glutamine, and 1x antibiotic-antimycotic solution (Thermo Fisher Scientific), at 37 °C, 5 % CO_2_, as described previously [30].

### Hi-C sample preparation using the original protocol

We have made modifications to a protocol introduced in previous literature [23, 31] (Fig. 1B). The full version of the modified ‘inexpensive and controllable Hi-C (iconHi-C)’ protocol is described in Supplementary Protocol S1.

### Hi-C sample preparation using commercial kits

The Proximo Hi-C kit (Phase Genomics) which employs the restriction enzyme Sau3A1 and transposase-based library preparation [32] (Fig. 1B) was used for preparing a library from the 50 mg softshell turtle liver following its official ver. 1.0 animal protocol (Library g in Fig. 7A) and a library from the 10 mg liver amplified with a reduced number of PCR cycles based on a preliminary real-time qPCR using an aliquot (Library h; see [25] for the detail of the pre-determination of optimal PCR cycles). The Arima Hi-C kit (Arima Genomics) which employs a restriction enzyme cocktail (Fig. 1B) was used in conjunction with the KAPA Hyper Prep Kit (KAPA Biosystems), protocol ver. A160108 v00, to prepare a library using the softshell turtle liver, following its official animal vertebrate tissue protocol (ver. A160107 v00) (Library f) and a library with an additional step of T4 DNA polymerase treatment for reducing ‘dangling end’ reads (Library e). This additional treatment is detailed in Step 8.2 (for DpnII-digested samples) in Supplementary Protocol S1.

### DNA sequencing

Small-scale sequencing for library QC was performed in-house to obtain 127 nt-long paired-end reads on an Illumina HiSeq 1500 in the Rapid Run Mode. Large-scale sequencing for Hi-C scaffolding was performed to obtain 151 nt-long paired-end reads on an Illumina HiSeq X. The obtained reads were subjected to quality control with FastQC ver. 0.11.5 (https://www.bioinformatics.babraham.ac.uk/projects/fastqc/), and low-quality regions and adapter sequences in the reads were removed using Trim Galore ver. 0.4.5 (https://www.bioinformatics.babraham.ac.uk/projects/trim_galore/) with the parameters ‘-e 0.1 −q 30’.

### Post-sequencing quality control of Hi-C libraries

For post-sequencing library QC, one million trimmed read pairs for each Hi-C library were sampled using the ‘subseq’ function of the program seqtk ver. 1.2-r94 (https://github.com/lh3/seqtk). The resultant sets of read pairs were processed using HiC-Pro ver. 2.11.1 [22] with bowtie2 ver. 2.3.4.1 [33] to evaluate the insert structure and mapping status onto the softshell turtle genome assembly PelSin_1.0 (GCF_000230535.1) or human genome assembly hg19. This resulted in the categorization between valid interaction pairs and invalid pairs, and the latter is divided into ‘dangling end’, ‘religation’, ‘self circle’, and ‘single-end’ (Fig. 4). To process the read pairs derived from the libraries prepared using either HindIII or DpnII (Sau3AI) with the iconHi-C protocol (Library a–d) and the Phase Genomics Proximo Hi-C kit (Library g and h), the restriction fragment file required by HiC-Pro was prepared according to the script ‘digest_genome.py’ provided with HiC-Pro. To process the reads derived from the Arima Hi-C kit (Library e and f), all restriction sites (‘GATC’ and ‘GANTC’) were inserted into the script. In addition, the nucleotide sequences of all possible ligated sites generated by restriction enzymes were included in a configuration file of HiC-Pro. The details and the sample code are included in Supplementary Protocol S2.

### Computation for Hi-C scaffolding

In order to control our comparison with intended input data sizes, certain numbers of trimmed read pairs were sampled for each library with seqtk as described above. Scaffolding was processed with the following methods employing two program pipelines, 3d-dna and SALSA2.

Scaffolding with the program 3d-dna was preceded by Hi-C read mapping onto the genome with Juicer ver. 20180805 [34] using the default parameters with BWA ver.0.7.17-r1188 [35]. The restriction fragment file required by Juicer was prepared by the script ‘generate_site_positions.py’ provided with Juicer or our original script compatible with multiple restriction enzymes to convert the restriction fragment file of HiC-Pro to the format required by Juicer (Supplementary Protocol S2). Scaffolding with 3d-dna ver. 20180929 was performed with variable parameters (see Fig. 9A).

Scaffolding with the program SALSA2 using Hi-C reads was preceded by Hi-C read pair processing with the Arima mapping pipeline ver. 20181207 (https://github.com/ArimaGenomics/mapping_pipeline) together with BWA, SAMtools ver. 1.8-21-gf6f50ac [36] and Picard ver. 2.18.12 (https://github.com/broadinstitute/picard). The mapping result in the binary alignment map (bam) format was converted into a BED file by bamToBed of Bedtools ver. 2.26.0 [37], whose output was used as an input of scaffolding using SALSA2 ver. 20181212 with the default parameters.

### Completeness assessment of Hi-C scaffolds

gVolante ver. 1.2.1 [28] was used to perform an assessment of sequence length distribution and gene space completeness based on the coverage of one-to-one reference orthologs with BUSCO v2/v3 employing the one-to-one ortholog set ‘Tetrapoda’ supplied with BUSCO [38]. For the assessment, no threshold of cut-off length was set.

### Continuity assessment with RNA-seq read mapping

Paired-end reads obtained by RNA-seq of softshell turtle embryos at multiple stages were downloaded from NCBI SRA (DRX001576) and were assembled with the program Trinity ver. 2.7.0 [39] with the default parameters. The assembled transcript sequences were mapped with pblat [40] to the Hi-C scaffold sequences, and the output was assessed with isoblat ver. 0.31 [41].

### Comparison with chromosome FISH results

Cytogenetic validation of Hi-C scaffolding results was performed by comparing the gene locations on the scaffold sequences with those in preexisting chromosome FISH data for 162 protein-coding genes [17–19]. The nucleotide exonic sequences for those 162 genes retrieved from GenBank were aligned with Hi-C scaffold sequences using BLAT ver. 36×2 [42], and their positions and orientation along the Hi-C scaffold sequences were analyzed.

### Additional files

Supplementary Figure S1. Quality control of the Hi-C libraries.

Supplementary Figure S2. Structural analysis of the possibly overassembled scaffold in Assembly #8

Supplementary Figure S3. Results of quality controls before sequencing.

Supplementary Table S1. Statistics of Chinese softshell turtle draft genome assembly before Hi-C.

Supplementary Table S2. HiC-Pro results of the human GM12878 HindIII Hi-C library with reduced reads

Supplementary Table S3. Quality control of Hi-C libraries.

Supplementary Table S4. Scaffolding results with variable input data and computational parameters

Supplementary Table S5. Mapping results of assembled transcript sequences onto Hi-C scaffolds

Supplementary Table S6. HiC-Pro results of the softshell turtle liver DpnII library (Library d) with reduced reads

Supplementary Table S7. Quality control of the human GM12878 Hi-C libraries

Supplementary Protocol S1. Protocol of iconHi-C

Supplementary Protocol S2. Computational protocol to support multiple enzymes

## Supporting information

Supplementary Tables & Protocols

## Abbreviations

PCR: polymerase chain reaction
FISH: fluorescence *in situ* hybridization
BUSCO: benchmarking universal single-copy orthologs
NCBI: National Center for Biotechnology Information
NGS: next generation DNA sequencing

## Acknowledgements

The authors acknowledge Naoki Irie, Juan Pascual Anaya and Tatsuya Hirasawa in Laboratory for Evolutionary Morphology, RIKEN BDR for suggestions for sampling, Rawin Poonperm for comments and discussion on the iconHi-C protocol, Olga Dudchenko, Erez Lieberman-Aiden, Arang Rhie, Sergey Koren, and Jay Ghurye for their technical suggestions for sample preparation and computation, Yoshinobu Uno for guidance to cytogenetic data interpretation, and Anthony Schmitt of Arima Genomics and Stephen Eacker of Phase Genomics for providing information about the Hi-C kits. They also thank the other members of Laboratory for Phyloinformatics and Laboratory for Developmental Epigenetics in RIKEN BDR for technical support and discussion.

## Author contributions

S.K., I.H., H.M., and M.K. conceived the study. M.K. and K.T. performed laboratory works, and O.N. performed bioinformatic analysis. M.K., O.N., and H.M. analyzed the data. S.K., M.K., and O.N. drafted the manuscript. All authors contributed to the finalization of the manuscript.

---

**Supplementary Figure S1:**
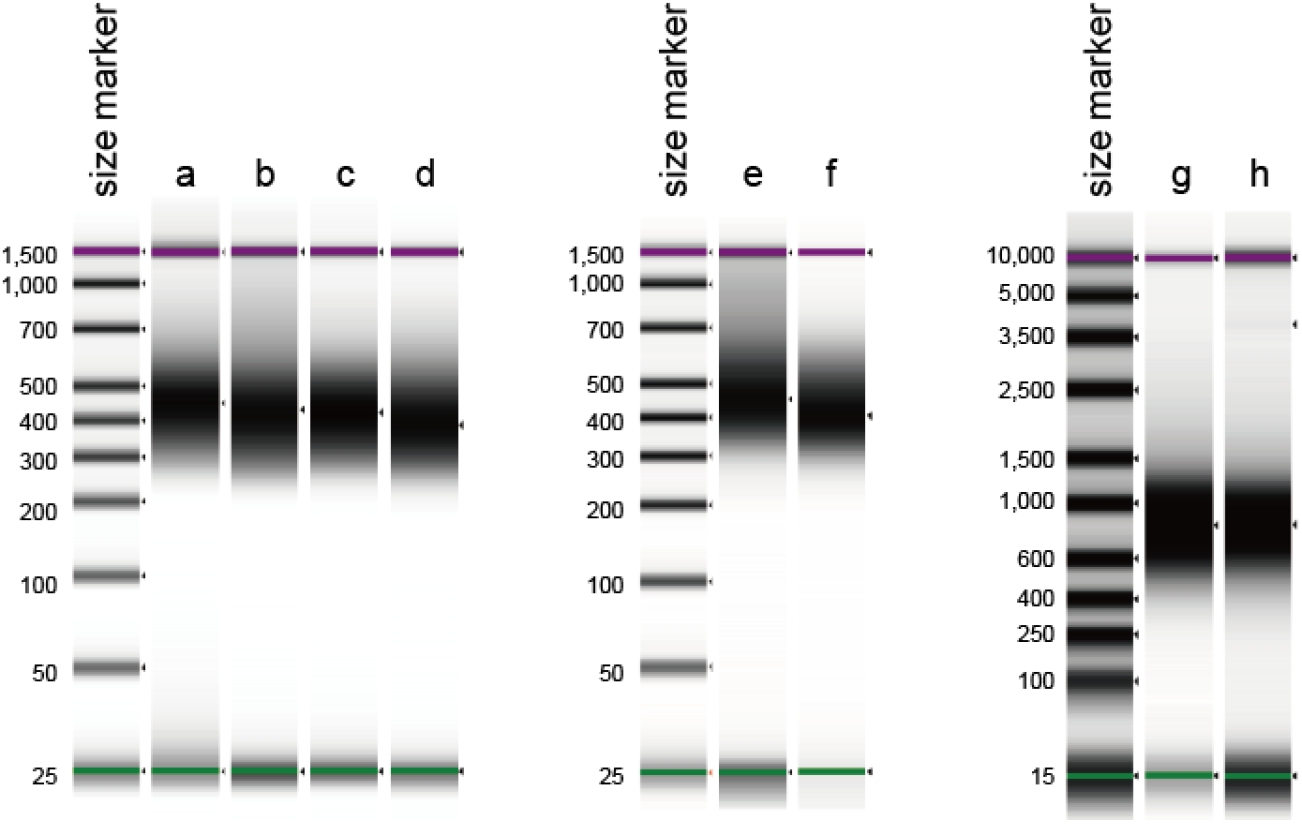
DNA size distribution of the softshell turtle Hi-C libraries. Size distribution of the libraries was analyzed by Agilent 4200 TapeStation using the High Sensitivity D1000 kit for Library a-f and the High Sensitivity D5000 kit for Library g and h.

**Supplementary Figure S2:**
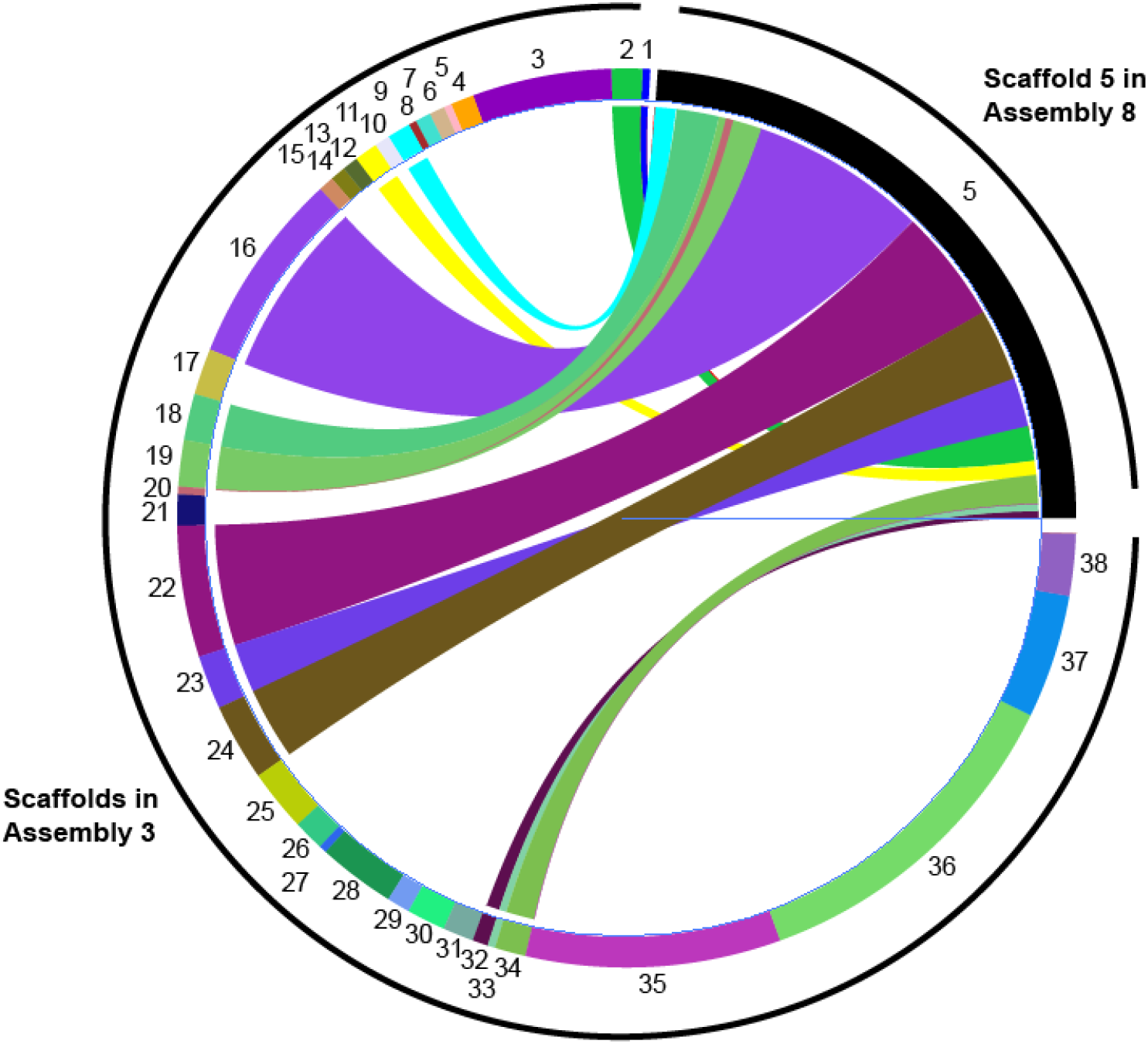
Structural analysis of the possibly overassembled scaffold in Assembly 8. This figure shows the nucleotide sequence-level correspondence of the whole sequence of the scaffold 5 of Assembly 8 to 14 scaffolds of Assembly 3. Note that the scaffold 5 of Assembly 8 accounts for approximately one-third of the estimated genome size, and that some of the scaffolds of Assembly 3 in the figure have multiple high-similarity regions in the scaffold 5 of Assembly 8.

**Supplementary Figure S3:**
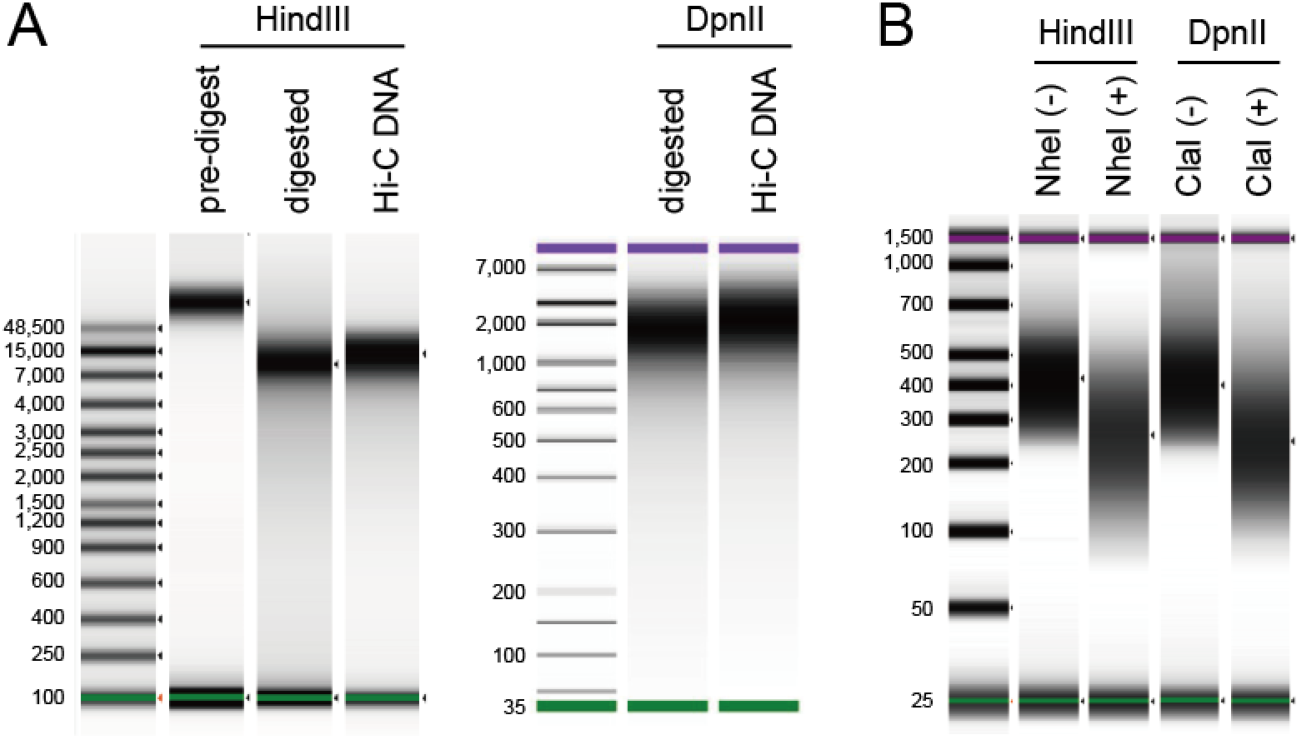
Pre-sequencing quality control of softshell turtle blood Hi-C libraries (Library a and b). (A) Quality control of Hi-C DNAs (QC1). Hi-C DNA was prepared from the Chinese softshell turtle blood by HindIII or DpnII digestion (see Fig. 7A for the detail). (B) Quality control of Hi-C libraries (QC2). The prepared softshell turtle blood library employing HindIII was digested by NheI, and the one employing DpnII was digested by ClaI (see Fig. 3 for the technical principle).

## References

1. Rowley MJ and Corces VG. Organizational principles of 3D genome architecture. Nature Reviews Genetics. 2018;19 12:789–800. doi:10.1038/s41576-018-0060-8.

2. Lieberman-Aiden E, van Berkum NL, Williams L, Imakaev M, Ragoczy T, Telling A, et al. Comprehensive Mapping of Long-Range Interactions Reveals Folding Principles of the Human Genome. Science. 2009;326 5950:289–93. doi:10.1126/science.1181369.

3. Rao Suhas SP, Huntley Miriam H, Durand Neva C, Stamenova Elena K, Bochkov Ivan D, Robinson James T, et al. A 3D Map of the Human Genome at Kilobase Resolution Reveals Principles of Chromatin Looping. Cell. 2014;159 7:1665–80. doi:10.1016/j.cell.2014.11.021.

4. Burton JN, Adey A, Patwardhan RP, Qiu R, Kitzman JO and Shendure J. Chromosome-scale scaffolding of de novo genome assemblies based on chromatin interactions. Nature Biotechnology. 2013;31:1119. doi:10.1038/nbt.2727.

5. Sedlazeck FJ, Lee H, Darby CA and Schatz MC. Piercing the dark matter: bioinformatics of long-range sequencing and mapping. Nature Reviews Genetics. 2018;19 6:329–46. doi:10.1038/s41576-018-0003-4.

6. Bickhart DM, Rosen BD, Koren S, Sayre BL, Hastie AR, Chan S, et al. Single-molecule sequencing and chromatin conformation capture enable de novo reference assembly of the domestic goat genome. Nature Genetics. 2017;49:643. doi:10.1038/ng.3802.

7. Putnam NH, O’Connell BL, Stites JC, Rice BJ, Blanchette M, Calef R, et al. Chromosome-scale shotgun assembly using an in vitro method for long-range linkage. Genome Research. 2016; doi:10.1101/gr.193474.115.

8. Ghurye J, Pop M, Koren S, Bickhart D and Chin C-S. Scaffolding of long read assemblies using long range contact information. BMC Genomics. 2017;18 1:527. doi:10.1186/s12864-017-3879-z.

9. Ghurye J, Rhie A, Walenz BP, Schmitt A, Selvaraj S, Pop M, et al. Integrating Hi-C links with assembly graphs for chromosome-scale assembly. bioRxiv. 2018:261149. doi:10.1101/261149.

10. Dudchenko O, Batra SS, Omer AD, Nyquist SK, Hoeger M, Durand NC, et al. De novo assembly of the Aedes aegypti genome using Hi-C yields chromosome-length scaffolds. Science. 2017;356 6333:92–5. doi:10.1126/science.aal3327.

11. Lewin HA, Robinson GE, Kress WJ, Baker WJ, Coddington J, Crandall KA, et al. Earth BioGenome Project: Sequencing life for the future of life. Proceedings of the National Academy of Sciences of the United States of America. 2018;115 17:4325–33. doi:10.1073/pnas.1720115115.

12. Koepfli KP, Paten B and O’Brien SJ. The Genome 10K Project: a way forward. Annual review of animal biosciences. 2015;3:57–111. doi:10.1146/annurev-animal-090414-014900.

13. Editorial. A reference standard for genome biology. Nature Biotechnology. 2018;36:1121. doi:10.1038/nbt.4318.

14. Dudchenko O, Shamim MS, Batra SS, Durand NC, Musial NT, Mostofa R, et al. The Juicebox Assembly Tools module facilitates de novo assembly of mammalian genomes with chromosome-length scaffolds for under $1000. bioRxiv. 2018:254797. doi:10.1101/254797.

15. Belaghzal H, Dekker J and Gibcus JH. Hi-C 2.0: An optimized Hi-C procedure for high-resolution genome-wide mapping of chromosome conformation. Methods (San Diego, Calif). 2017;123:56–65. doi:10.1016/j.ymeth.2017.04.004.

16. Kuratani S, Kuraku S and Nagashima H. Evolutionary developmental perspective for the origin of turtles: the folding theory for the shell based on the developmental nature of the carapacial ridge. Evolution & Development. 2011;13 1:1–14. doi:10.1111/j.1525-142X.2010.00451.x.

17. Matsuda Y, Umehara C, Nishida-Tarui H, Kuroiwa A, Yamada K, Isobe T, et al. Highly conserved linkage homology between birds and turtles: bird and turtle chromosomes are precise counterparts of each other. Chromosome research : an international journal on the molecular, supramolecular and evolutionary aspects of chromosome biology. 2005;13 6:601–15. doi:10.1007/s10577-005-0986-5.

18. Kuraku S, Ishijima J, Nishida-Umehara C, Agata K, Kuratani S and Matsuda Y. cDNA-based gene mapping and GC3 profiling in the soft-shelled turtle suggest a chromosomal size-dependent GC bias shared by sauropsids. Chromosome research : an international journal on the molecular, supramolecular and evolutionary aspects of chromosome biology. 2006;14 2:187–202. doi:10.1007/s10577-006-1035-8.

19. Uno Y, Nishida C, Tarui H, Ishishita S, Takagi C, Nishimura O, et al. Inference of the protokaryotypes of amniotes and tetrapods and the evolutionary processes of microchromosomes from comparative gene mapping. PloS one. 2012;7 12:e53027. doi:10.1371/journal.pone.0053027.

20. Wang Z, Pascual-Anaya J, Zadissa A, Li W, Niimura Y, Huang Z, et al. The draft genomes of soft-shell turtle and green sea turtle yield insights into the development and evolution of the turtle-specific body plan. Nature Genetics. 2013;45:701. doi:10.1038/ng.2615.

21. Belton JM, McCord RP, Gibcus JH, Naumova N, Zhan Y and Dekker J. Hi-C: a comprehensive technique to capture the conformation of genomes. Methods. 2012;58 3:268–76. doi:10.1016/j.ymeth.2012.05.001.

22. Servant N, Varoquaux N, Lajoie BR, Viara E, Chen CJ, Vert JP, et al. HiC-Pro: an optimized and flexible pipeline for Hi-C data processing. Genome Biol. 2015;16:259. doi:10.1186/s13059-015-0831-x.

23. Sofueva S, Yaffe E, Chan WC, Georgopoulou D, Vietri Rudan M, Mira-Bontenbal H, et al. Cohesin-mediated interactions organize chromosomal domain architecture. The EMBO journal. 2013;32 24:3119–29. doi:10.1038/emboj.2013.237.

24. Mayjonade B, Gouzy J, Donnadieu C, Pouilly N, Marande W, Callot C, et al. Extraction of high-molecular-weight genomic DNA for long-read sequencing of single molecules. BioTechniques. 2016;61 4:203–5. doi:10.2144/000114460.

25. Tanegashima C, Nishimura O, Motone F, Tatsumi K, Kadota M and Kuraku S. Embryonic transcriptome sequencing of the ocellate spot skate Okamejei kenojei. Scientific data. 2018;5:180200. doi:10.1038/sdata.2018.200.

26. Botero-Castro F, Figuet E, Tilak MK, Nabholz B and Galtier N. Avian Genomes Revisited: Hidden Genes Uncovered and the Rates versus Traits Paradox in Birds. Molecular biology and evolution. 2017;34 12:3123–31. doi:10.1093/molbev/msx236.

27. Hotaling S and Kelley JL. The rising tide of high-quality genomic resources. Molecular Ecology Resources. 2019;19 3:567–9. doi:10.1111/1755-0998.12964.

28. Nishimura O, Hara Y and Kuraku S. gVolante for standardizing completeness assessment of genome and transcriptome assemblies. Bioinformatics (Oxford, England). 2017;33 22:3635–7. doi:10.1093/bioinformatics/btx445.

29. Roach MJ, Schmidt SA and Borneman AR. Purge Haplotigs: allelic contig reassignment for third-gen diploid genome assemblies. BMC Bioinformatics. 2018;19 1:460. doi:10.1186/s12859-018-2485-7.

30. Kadota M, Hara Y, Tanaka K, Takagi W, Tanegashima C, Nishimura O, et al. CTCF binding landscape in jawless fish with reference to Hox cluster evolution. Scientific Reports. 2017;7 1:4957. doi:10.1038/s41598-017-04506-x.

31. Miura H, Takahashi S, Poonperm R, Tanigawa A, Takebayashi S and Hiratani I. Spatiotemporal developmental dynamics of chromosome organization revealed by single-cell DNA replication profiling. in press.

32. Adey A, Morrison HG, Asan, Xun X, Kitzman JO, Turner EH, et al. Rapid, low-input, low-bias construction of shotgun fragment libraries by high-density in vitro transposition. Genome Biology. 2010;11 12:R119. doi:10.1186/gb-2010-11-12-r119.

33. Langmead B and Salzberg SL. Fast gapped-read alignment with Bowtie 2. Nature Methods. 2012;9:357. doi:10.1038/nmeth.1923.

34. Durand NC, Shamim MS, Machol I, Rao SS, Huntley MH, Lander ES, et al. Juicer Provides a One-Click System for Analyzing Loop-Resolution Hi-C Experiments. Cell systems. 2016;3 1:95–8. doi:10.1016/j.cels.2016.07.002.

35. Li H and Durbin R. Fast and accurate short read alignment with Burrows-Wheeler transform. Bioinformatics (Oxford, England). 2009;25 14:1754–60. doi:10.1093/bioinformatics/btp324.

36. Li H. A statistical framework for SNP calling, mutation discovery, association mapping and population genetical parameter estimation from sequencing data. Bioinformatics (Oxford, England). 2011;27 21:2987–93. doi:10.1093/bioinformatics/btr509.

37. Quinlan AR and Hall IM. BEDTools: a flexible suite of utilities for comparing genomic features. Bioinformatics (Oxford, England). 2010;26 6:841–2. doi:10.1093/bioinformatics/btq033.

38. Simao FA, Waterhouse RM, Ioannidis P, Kriventseva EV and Zdobnov EM. BUSCO: assessing genome assembly and annotation completeness with single-copy orthologs. Bioinformatics (Oxford, England). 2015;31 19:3210–2. doi:10.1093/bioinformatics/btv351.

39. Grabherr MG, Haas BJ, Yassour M, Levin JZ, Thompson DA, Amit I, et al. Full-length transcriptome assembly from RNA-Seq data without a reference genome. Nat Biotechnol. 2011;29 7:644–52. doi:10.1038/nbt.1883.

40. Wang M and Kong L. pblat: a multithread blat algorithm speeding up aligning sequences to genomes. BMC Bioinformatics. 2019;20 1:28. doi:10.1186/s12859-019-2597-8.

41. Ryan JF. Baa.pl: A tool to evaluate de novo genome assemblies with RNA transcripts. arXiv e-prints. 2013.

42. Kent WJ. BLAT--the BLAST-like alignment tool. Genome Res. 2002;12 4:656–64. doi:10.1101/gr.229202.

43. Imakaev M, Fudenberg G, McCord RP, Naumova N, Goloborodko A, Lajoie BR, et al. Iterative correction of Hi-C data reveals hallmarks of chromosome organization. Nature Methods. 2012;9:999. doi:10.1038/nmeth.2148.

