## Supplementary Tables & Protocols for "Multifaceted Hi-C benchmarking: what makes a difference in chromosome-scale genome scaffolding?"

**Supplementary Table S1:** Statistics of Chinese softshell turtle draft genome assembly before Hi-C.

| **Sequence length statistics** | |
|---|---|
| Number of scaffolds | 19,903 |
| Total length of scaffolds | 2.20 Gb |
| Maximum scaffold length | 16.02 Mb |
| Minimum scaffold length | 200 b |
| Number of sequences longer than 1K nt | 5,962 |
| Number of sequences longer than 10K nt | 3,215 |
| Number of sequences longer than 100K nt | 1,491 |
| Number of sequences longer than 1M nt | 558 |
| Number of sequences longer than 10M nt | 14 |
| Mean scaffold length | 110.7 Kb |
| N50 scaffold length | 3.35 Mb |
| Overall GC-content | 44.41% |
| Overall N-content | 4.35% |
| Sum length proportion of sequences longer than 1M nt | 81.00% |
| Sum length proportion of sequences longer than 10M nt | 7.40% |

| **Completeness of gene space inferred by BUSCO** | |
|---|---|
| Total number of reference orthologs queried (Tetrapoda) | 3,950 |
| Number of reference orthologs detected | |
| Complete | 3,624 |
| Complete + fragmented | 3,819 |
| Number of missing orthologs | 131 |
| Average number of copies per ortholog | 1.01 |
| Detected orthologs that have more than one copy | 0.72% |

**Supplementary Table S2:** HiC-Pro results of the human GM12878 HindIII Hi-C library with reduced reads

**A. Read alignment category**

| | Proportion of reads | | | | | | |
|---|---|---|---|---|---|---|---|
| Number of input read pairs | 500K | 1M | 5M | 10M | 50M | 100M | 200M |
| Unique paired alignments | 71.0% | 71.0% | 70.9% | 71.0% | 71.0% | 71.0% | 71.0% |
| Unmapped pairs | 3.2% | 3.2% | 3.2% | 3.2% | 3.2% | 3.2% | 3.2% |
| Low quality pairs | 0.0% | 0.0% | 0.0% | 0.0% | 0.0% | 0.0% | 0.0% |
| Multiple pairs alignments | 15.3% | 15.3% | 15.3% | 15.3% | 15.3% | 15.3% | 15.3% |
| Pairs with singleton | 10.5% | 10.5% | 10.5% | 10.5% | 10.5% | 10.5% | 10.5% |
| Low quality singleton | 0.0% | 0.0% | 0.0% | 0.0% | 0.0% | 0.0% | 0.0% |
| Unique singleton alignments | 0.0% | 0.0% | 0.0% | 0.0% | 0.0% | 0.0% | 0.0% |
| Multiple singleton alignments | 0.0% | 0.0% | 0.0% | 0.0% | 0.0% | 0.0% | 0.0% |
| Reported pairs | 71.0% | 71.0% | 70.9% | 71.0% | 71.0% | 71.0% | 71.0% |

**B. Read pair category**

| | Proportion of read pairs | | | | | | |
|---|---|---|---|---|---|---|---|
| Number of input read pairs | 500K | 1M | 5M | 10M | 50M | 100M | 200M |
| Valid interaction pairs | 65.1% | 65.1% | 65.1% | 65.1% | 65.1% | 65.1% | 65.1% |
| Valid interaction pairs (forward-forward) | 16.2% | 16.2% | 16.2% | 16.2% | 16.2% | 16.2% | 16.2% |
| Valid interaction pairs (reverse-reverse) | 16.1% | 16.2% | 16.2% | 16.2% | 16.2% | 16.2% | 16.2% |
| Valid interaction pairs (reverse-forward) | 15.8% | 15.8% | 15.7% | 15.7% | 15.7% | 15.7% | 15.7% |
| Valid interaction pairs (forward-reverse) | 17.0% | 17.0% | 16.9% | 16.9% | 16.9% | 16.9% | 16.9% |
| Dangling end pairs | 2.8% | 2.9% | 2.9% | 2.9% | 2.9% | 2.9% | 2.9% |
| Religation pairs | 2.6% | 2.5% | 2.6% | 2.6% | 2.6% | 2.6% | 2.6% |
| Self circle pairs | 0.5% | 0.4% | 0.4% | 0.4% | 0.4% | 0.4% | 0.4% |
| Single-end pairs | 0.0% | 0.0% | 0.0% | 0.0% | 0.0% | 0.0% | 0.0% |
| Filtered pairs | 0.0% | 0.0% | 0.0% | 0.0% | 0.0% | 0.0% | 0.0% |
| Dumped pairs | 0.0% | 0.0% | 0.0% | 0.0% | 0.0% | 0.0% | 0.0% |

**C. Duplicates and contact ranges**

| | Proportion of read pairs | | | | | | |
|---|---|---|---|---|---|---|---|
| Number of input read pairs | 500K | 1M | 5M | 10M | 50M | 100M | 200M |
| Valid interaction | 65.1% | 65.1% | 65.1% | 65.1% | 65.1% | 65.1% | 65.1% |
| Valid interaction (remove duplicates) | 65.1% | 65.0% | 64.8% | 64.5% | 62.3% | 59.8% | 55.2% |
| Trans interaction | 12.0% | 12.0% | 12.0% | 11.9% | 11.5% | 11.1% | 10.2% |
| Cis interaction | 53.1% | 53.1% | 52.8% | 52.6% | 50.8% | 48.7% | 45.0% |
| Cis short-range interaction (<20kb) | 7.2% | 7.2% | 7.1% | 7.1% | 6.8% | 6.5% | 6.0% |
| Cis long-range interaction (>20kb) | 45.9% | 45.9% | 45.7% | 45.5% | 44.0% | 42.2% | 38.9% |

Note that the *cis* / *trans* judgement based on fragmentary sequences might be unreliable because insufficient continuity increases the trans fraction and instead decreases the cis fraction.

**Supplementary Table S3:** Quality control of Hi-C libraries.

| Library preparation condition | Library ID | | | | | | | |
|---|---|---|---|---|---|---|---|---|
| | **a** | **b** | **c** | **d** | **e** | **f** | **g** | **h** |
| Tissue type | Blood | | Liver | | | | | |
| Amount of cells/tissue or DNA used for the Hi-C reaction | 2 x $10^6$ cells | | Tissue estimated to contain 3 µg DNA (based on the iconHi-C protocol: Additional file Protocol Step 3) | | 10 mg tissue (estimated to contain 4.23 µg DNA based on the input material determination protocol of the Arima-HiC kit) | | 50 mg tissue (estimated to contain 110.0 ng DNA based on the DNA QC step performed after restriction digestion) | 10 mg tissue (estimated to contain 66.3 ng DNA based on the DNA QC step performed after restriction digestion) |
| Restriction enzyme | HindIII | DpnII | HindIII | DpnII | Arima cocktail | | Sau3AI | |
| Hi-C DNA amount for lib prep (µg) | 2 | 1 | 2 | 1 | 1 | 1 | N/A | |
| Number of PCR cycles | 6 | 8 | 6 | 5 | 8 | 9 | 15 | 11 |
| Average library size (bp) | 488 | 493 | 456 | 444 | 558 | 454 | 892 (1,052 bp before size selection) | 882 (1,100 bp before size selection) |
| Library yield (ng) | 112.2 | 129.6 | 140.4 | 172.2 | 160.0 | 1,180.0 | 232.0 (1,008 ng before size selection) | 32.8 (88 ng before size selection) |
| Additional treatment (divergences from the original protocol) | N/A | | | | Treatment of the Hi-C DNA with T4 DNA polymerase | N/A | Size selection of the Hi-C library with AMPure XP | Size selection of the Hi-C library with AMPure XP; Reduced number of PCR cycles |
| **Read alignment category** | **Proportion of read pairs (%)** | | | | | | | |
| Unique paired alignments | 57.1 | 55.9 | 54.2 | 60.1 | 65.2 | 69.3 | 68.2 | 62.7 |
| Unmapped pairs | 5.7 | 3.8 | 8.0 | 2.5 | 2.3 | 1.8 | 4.0 | 5.1 |
| Low quality pairs | 0.0 | 0.0 | 0.0 | 0.0 | 0.0 | 0.0 | 0.0 | 0.0 |
| Multiple pairs alignments | 16.1 | 19.4 | 14.4 | 20.6 | 15.1 | 14.4 | 9.1 | 8.8 |
| Pairs with singleton | 21.2 | 20.9 | 23.4 | 16.8 | 17.4 | 14.5 | 18.7 | 23.4 |
| Low quality singleton | 0.0 | 0.0 | 0.0 | 0.0 | 0.0 | 0.0 | 0.0 | 0.0 |
| Unique singleton alignments | 0.0 | 0.0 | 0.0 | 0.0 | 0.0 | 0.0 | 0.0 | 0.0 |
| Multiple singleton alignments | 0.0 | 0.0 | 0.0 | 0.0 | 0.0 | 0.0 | 0.0 | 0.0 |
| Reported pairs | 57.1 | 55.9 | 54.2 | 60.1 | 65.2 | 69.3 | 68.2 | 62.7 |
| **Read pair category** | **Proportion of read pairs (%)** | | | | | | | |
| Valid interaction pairs | 52.4 | 52.1 | 48.1 | 58.8 | 63.6 | 64.7 | 36.5 | 33.5 |
| Dangling end pairs | 2.2 | 2.8 | 3.8 | 0.7 | 0.5 | 2.5 | 20.4 | 20.1 |
| Religation pairs | 2.1 | 0.7 | 1.9 | 0.5 | 1.1 | 2.1 | 11.1 | 8.9 |
| Self circle pairs | 0.4 | 0.3 | 0.3 | 0.1 | 0.0 | 0.0 | 0.1 | 0.1 |
| Single-end pairs | 0.0 | 0.0 | 0.0 | 0.0 | 0.0 | 0.0 | 0.0 | 0.0 |
| Filtered pairs | 0.0 | 0.0 | 0.0 | 0.0 | 0.0 | 0.0 | 0.0 | 0.0 |
| Dumped pairs | 0.1 | 0.0 | 0.0 | 0.0 | 0.0 | 0.0 | 0.0 | 0.0 |

See Fig. 4 for the detail of read pair categorization.

**Supplementary Table S4:** Scaffolding results with variable input data and computational parameters

| Assembly ID | Sample preparation condition | | | | Scaffolding condition | | | | Basic sequence compositions | | | | | | | | | | | | | Gene space completeness assessment by BUSCO referring to 3,950 Tetrapoda BUSCOs | | | | | | | | |
|---|---|---|---|---|---|---|---|---|---|---|---|---|---|---|---|---|---|---|---|---|---|---|---|---|---|---|---|---|---|---|
| | Library ID | Preparation method | Tissue(s) | Restriction enzyme(s) | Scaffolding program | Smaller input sequence cutoff length* | Numbers of iterative corrections** | Number of input read pairs | Number of scaffolds | Maximum scaffold length | Number of sequences longer than 1K nt | Number of sequences longer than 10K nt | Number of sequences longer than 100K nt | Number of sequences longer than 1M nt | Number of sequences longer than 10M nt | Mean scaffold length | Median scaffold length | N50 scaffold length | Overall N-content (%) | Sum proportion of sequences longer than 1M nt (%) | Sum proportion of sequences longer than 10M nt (%) | Complete + Fragmented BUSCOs (C + F) | Complete and single-copy BUSCOs (S) | Complete and duplicated BUSCOs (D) | Fragmented BUSCOs (F) | Missing BUSCOs (M) | Complete BUSCOs (C) | Average number of copies per ortholog | Detected orthologs that have more than one copy (%) | Scores in percentages |
| Assembly before Hi-C | | | | | | | | | 19,903 | 16,024,077 | 5,962 | 3,215 | 1491 | 558 | 14 | 110,660 | 498 | 3,350,749 | 4.35 | 81.0 | 7.4 | 3,819 | 3,598 | 26 | 195 | 131 | 3,624 | 1.01 | 0.72 | C:91.8%[S:91.1%,D:0.7%],F:4.9%,M:3.3%,n:3950 |
| **1** | c | iconHi-C | liver | HindIII | 3d-dna | 3d-dna defaults | | 200,000,000 | 19,824 | 347,378,596 | 5,807 | 2,283 | 72 | 46 | 34 | 111,169 | 496 | 128,648,452 | 4.41 | 96.3 | 93.5 | 3,807 | 3,566 | 30 | 211 | 143 | 3,596 | 1.01 | 0.83 | C:91.1%[S:90.3%,D:0.8%],F:5.3%,M:3.6%,n:3950 |
| **2** | a | | blood | | | | | | 20,899 | 151,823,619 | 6,804 | 3,203 | 197 | 91 | 47 | 105,456 | 529 | 40,769,607 | 4.41 | 94.0 | 83.3 | 3,811 | 3,547 | 27 | 237 | 139 | 3,574 | 1.01 | 0.76 | C:90.5%[S:89.8%,D:0.7%],F:6.0%,M:3.5%,n:3950 |
| **3** | d | | liver | DpnII | | | | | 18,013 | 352,879,942 | 4,005 | 740 | 40 | 38 | 31 | 122,341 | 439 | 130,239,026 | 4.41 | 98.7 | 96.9 | 3,821 | 3,619 | 27 | 175 | 129 | 3,646 | 1.01 | 0.74 | C:92.3%[S:91.6%,D:0.7%],F:4.4%,M:3.3%,n:3950 |
| **4** | b | | blood | | | | | | 18,215 | 140,927,669 | 4,161 | 821 | 117 | 108 | 62 | 120,984 | 446 | 31,364,868 | 4.41 | 98.5 | 87.8 | 3,820 | 3,604 | 31 | 185 | 130 | 3,635 | 1.01 | 0.85 | C:92.0%[S:91.2%,D:0.8%],F:4.7%,M:3.3%,n:3950 |
| **5** | c | | liver | HindIII | SALSA2 | SALSA2 defaults | | | 18,674 | 102,051,213 | 4,762 | 2,051 | 577 | 227 | 53 | 117,978 | 462 | 18,089,892 | 4.38 | 92.0 | 63.0 | 3,826 | 3,615 | 30 | 181 | 124 | 3,645 | 1.01 | 0.82 | C:92.3%[S:91.5%,D:0.8%],F:4.6%,M:3.1%,n:3950 |
| **6** | d | | | DpnII | | | | | 18,469 | 142,087,433 | 4,537 | 1,802 | 500 | 215 | 56 | 119,293 | 455 | 17,382,281 | 4.38 | 93.3 | 65.9 | 3,823 | 3,603 | 31 | 189 | 127 | 3,634 | 1.01 | 0.85 | C:92.0%[S:91.2%,D:0.8%],F:4.8%,M:3.2%,n:3950 |
| **7** | c+d | | liver | HindIII & DpnII | 3d-dna | 3d-dna defaults | | | 18,087 | 351,946,035 | 4,084 | 766 | 50 | 42 | 34 | 121,838 | 441 | 130,589,277 | 4.40 | 98.6 | 96.8 | 3,820 | 3,624 | 29 | 167 | 130 | 3,653 | 1.01 | 0.79 | C:92.4%[S:91.7%,D:0.7%],F:4.2%,M:3.4%,n:3950 |
| **8** | b+d | | blood & liver | DpnII | | | | | 17,937 | 703,313,584 | 3,922 | 690 | 18 | 15 | 15 | 122,863 | 436 | 303,431,936 | 4.41 | 98.7 | 98.7 | 3,809 | 3,611 | 25 | 173 | 141 | 3,636 | 1.01 | 0.69 | C:92.0%[S:91.4%,D:0.6%],F:4.4%,M:3.6%,n:3950 |
| **9** | e | Arima kit | liver | cocktail of A1 and A2 | 3d-dna | 3d-dna defaults | | | 17,756 | 352,772,394 | 3,730 | 670 | 47 | 43 | 35 | 124,117 | 429 | 130,552,621 | 4.41 | 98.8 | 97.4 | 3,822 | 3,616 | 30 | 176 | 128 | 3,646 | 1.01 | 0.82 | C:92.3%[S:91.5%,D:0.8%],F:4.5%,M:3.2%,n:3950 |
| **10** | e | | | | SALSA2 | SALSA2 defaults | | | 18,374 | 86,047,802 | 4,450 | 1,734 | 505 | 232 | 60 | 119,913 | 452 | 14,588,975 | 4.39 | 93.5 | 60.9 | 3,817 | 3,599 | 27 | 191 | 133 | 3,626 | 1.01 | 0.74 | C:91.8%[S:91.1%,D:0.7%],F:4.8%,M:3.4%,n:3950 |
| **11** | h | Phase kit | liver | Sau3AI | 3d-dna | 3d-dna defaults | | | 20,629 | 132,711,915 | 6,308 | 1,751 | 160 | 99 | 47 | 106,827 | 520 | 43,805,214 | 4.41 | 96.3 | 84.3 | 3,817 | 3,578 | 28 | 211 | 133 | 3,606 | 1.01 | 0.78 | C:91.3%[S:90.6%,D:0.7%],F:5.3%,M:3.4%,n:3950 |
| **12** | h | | | | SALSA2 | SALSA2 defaults | | | 18,690 | 65,503,740 | 4,900 | 2,241 | 745 | 293 | 55 | 117,877 | 467 | 9,689,575 | 4.38 | 90.2 | 49.6 | 3,822 | 3,591 | 28 | 203 | 128 | 3,619 | 1.01 | 0.77 | C:91.6%[S:90.9%,D:0.7%],F:5.1%,M:3.3%,n:3950 |
| **13** | d | iconHi-C | liver | DpnII | 3d-dna | -i 15000 | -r 2 | 200,000,000 | 18,013 | 352,879,942 | 4,005 | 740 | 40 | 38 | 31 | 122,341 | 439 | 130,239,026 | 4.41 | 98.7 | 96.9 | 3,821 | 3,619 | 27 | 175 | 129 | 3,646 | 1.01 | 0.74 | C:92.3%[S:91.6%,D:0.7%],F:4.4%,M:3.3%,n:3950 |
| **14** | d | | | | | -i 15000 | -r 4 | | 18,154 | 334,063,790 | 4,142 | 826 | 52 | 46 | 34 | 121,389 | 443 | 130,244,796 | 4.41 | 98.6 | 96.1 | 3,824 | 3,617 | 28 | 179 | 126 | 3,645 | 1.01 | 0.77 | C:92.3%[S:91.6%,D:0.7%],F:4.5%,M:3.2%,n:3950 |
| **15** | d | | | | | -i 15000 | -r 6 | | 18,199 | 352,652,404 | 4,188 | 845 | 35 | 29 | 22 | 121,087 | 445 | 156,729,449 | 4.40 | 98.5 | 97.3 | 3,814 | 3,619 | 25 | 170 | 136 | 3,644 | 1.01 | 0.69 | C:92.2%[S:91.6%,D:0.6%],F:4.3%,M:3.5%,n:3950 |
| **16** | d | | | | | -i 10000 | -r 2 | | 17,775 | 353,310,449 | 3,752 | 450 | 51 | 43 | 33 | 123,984 | 430 | 129,828,393 | 4.41 | 98.8 | 96.4 | 3,820 | 3,621 | 29 | 170 | 130 | 3,650 | 1.01 | 0.79 | C:92.4%[S:91.7%,D:0.7%],F:4.3%,M:3.3%,n:3950 |
| **17** | d | | | | | -i 5000 | -r 2 | | 17,424 | 353,773,630 | 3,404 | 391 | 54 | 44 | 34 | 126,493 | 419 | 129,814,843 | 4.42 | 98.9 | 96.6 | 3,817 | 3,621 | 28 | 168 | 133 | 3,649 | 1.01 | 0.77 | C:92.4%[S:91.7%,D:0.7%],F:4.3%,M:3.3%,n:3950 |
| **18** | d | | | | | -i 3000 | -r 2 | | 17,206 | 352,757,195 | 3,148 | 357 | 49 | 41 | 34 | 128,104 | 412 | 132,629,580 | 4.42 | 99.0 | 97.1 | 3,815 | 3,605 | 30 | 180 | 135 | 3,635 | 1.01 | 0.83 | C:92.1%[S:91.3%,D:0.8%],F:4.6%,M:3.3%,n:3950 |
| **19** | d | iconHi-C | liver | DpnII | 3d-dna | 3d-dna defaults | | 280,000,000 | 17,845 | 352,666,335 | 3,830 | 672 | 44 | 38 | 35 | 123,496 | 432 | 132,509,297 | 4.41 | 98.8 | 98.0 | 3,822 | 3,612 | 27 | 183 | 128 | 3,639 | 1.01 | 0.74 | C:92.1%[S:91.4%,D:0.7%],F:4.6%,M:3.3%,n:3950 |
| **20** | d | | | | | | | 160,000,000 | 18,129 | 352,585,997 | 4,128 | 800 | 60 | 54 | 33 | 121,556 | 442 | 130,809,328 | 4.40 | 98.6 | 93.7 | 3,817 | 3,607 | 29 | 181 | 133 | 3,636 | 1.01 | 0.80 | C:92.0%[S:91.3%,D:0.7%],F:4.6%,M:3.4%,n:3950 |
| **21** | d | | | | | | | 80,000,000 | 18,729 | 192,448,115 | 4,677 | 1,101 | 107 | 90 | 41 | 117,654 | 463 | 68,103,746 | 4.40 | 98.1 | 86.4 | 3,817 | 3,615 | 27 | 175 | 133 | 3,642 | 1.01 | 0.74 | C:92.2%[S:91.5%,D:0.7%],F:4.4%,M:3.4%,n:3950 |
| **22** | d | | | | | | | 20,000,000 | 33,115 | 42,618,151 | 17,006 | 7,765 | 503 | 208 | 65 | 66,681 | 1,146 | 11,576,731 | 4.60 | 85.2 | 56.0 | 3,730 | 3,322 | 30 | 378 | 220 | 3,352 | 1.01 | 0.89 | C:84.9%[S:84.1%,D:0.8%],F:9.6%,M:5.5%,n:3950 |
| **23** | d | | | | | | | 10,000,000 | 47,023 | 8,708,159 | 31,216 | 21,256 | 2475 | 367 | 0 | 47,049 | 7,307 | 616,243 | 4.78 | 35.6 | 0.0 | 3,583 | 2,875 | 28 | 680 | 367 | 2,903 | 1.01 | 0.96 | C:73.5%[S:72.8%,D:0.7%],F:17.2%,M:9.3%,n:3950 |

* Default parameters for 3d-dna and SALSA2 are 15000 and 1000, respectively.

** Default parameters for 3d-dna and SALSA2 are 2 and 3, respectively.

**Supplementary Table S5:** Mapping results of assembled transcript sequences onto Hi-C scaffolds

| Assembly ID* | Library ID** | Preparation method | Scaffolding program | Ratio of transcripts with a BLAT entry | Total % coverage of all positions | Number of transcripts mapping to a single contig/scaffold (ratio) | Average number of contigs/scaffolds per mapped transcript |
|---|---|---|---|---|---|---|---|
| 3 | d | iconHi-C | 3d-dna | 0.9877 | 0.9528 | 0.8790 | 1.1953 |
| 6 | d | iconHi-C | SALSA2 | 0.9877 | 0.9528 | 0.8722 | 1.2237 |
| 7 | c + d | iconHi-C | 3d-dna | 0.9877 | 0.9528 | 0.8784 | 1.1975 |
| 9 | e | Arima kit | 3d-dna | 0.9877 | 0.9528 | 0.8786 | 1.1981 |
| 11 | h | Phase kit | 3d-dna | 0.9878 | 0.9496 | 0.8700 | 1.2230 |

*See Fig. 9A for the detail.

**See Fig. 7A for the detail.

**Supplementary Table S6:** HiC-Pro results of the softshell turtle liver DpnII library (Library d) with reduced reads

**A. Read alignment category**

| | Proportion of read pairs | | | | | | |
|---|---|---|---|---|---|---|---|
| Number of input read pairs | 500K | 1M | 5M | 10M | 50M | 100M | 200M |
| Unique paired alignments | 59.4% | 59.4% | 59.3% | 59.3% | 59.3% | 59.3% | 59.3% |
| Unmapped pairs | 3.0% | 3.0% | 2.9% | 2.9% | 2.9% | 2.9% | 2.9% |
| Low quality pairs | 0.0% | 0.0% | 0.0% | 0.0% | 0.0% | 0.0% | 0.0% |
| Multiple pairs alignments | 20.1% | 20.0% | 20.1% | 20.1% | 20.1% | 20.1% | 20.1% |
| Pairs with singleton | 17.6% | 17.6% | 17.6% | 17.7% | 17.7% | 17.6% | 17.7% |
| Low quality singleton | 0.0% | 0.0% | 0.0% | 0.0% | 0.0% | 0.0% | 0.0% |
| Unique singleton alignments | 0.0% | 0.0% | 0.0% | 0.0% | 0.0% | 0.0% | 0.0% |
| Multiple singleton alignments | 0.0% | 0.0% | 0.0% | 0.0% | 0.0% | 0.0% | 0.0% |
| Reported pairs | 59.4% | 59.4% | 59.3% | 59.3% | 59.3% | 59.3% | 59.3% |

**B. Read pair category**

| | Proportion of read pairs | | | | | | |
|---|---|---|---|---|---|---|---|
| Number of input read pairs | 500K | 1M | 5M | 10M | 50M | 100M | 200M |
| Valid interaction pairs | 58.0% | 58.0% | 57.9% | 57.9% | 57.9% | 58.0% | 57.9% |
| Valid interaction pairs (forward-forward) | 14.3% | 14.5% | 14.4% | 14.4% | 14.5% | 14.5% | 14.5% |
| Valid interaction pairs (reverse-reverse) | 14.5% | 14.5% | 14.5% | 14.5% | 14.5% | 14.5% | 14.5% |
| Valid interaction pairs (reverse-forward) | 14.5% | 14.4% | 14.3% | 14.4% | 14.4% | 14.4% | 14.4% |
| Valid interaction pairs (forward-reverse) | 14.7% | 14.6% | 14.7% | 14.6% | 14.6% | 14.6% | 14.6% |
| Dangling end pairs | 0.8% | 0.8% | 0.8% | 0.8% | 0.8% | 0.8% | 0.8% |
| Religation pairs | 0.5% | 0.5% | 0.5% | 0.5% | 0.5% | 0.5% | 0.5% |
| Self circle pairs | 0.1% | 0.1% | 0.1% | 0.1% | 0.1% | 0.1% | 0.1% |
| Single-end pairs | 0.0% | 0.0% | 0.0% | 0.0% | 0.0% | 0.0% | 0.0% |
| Filtered pairs | 0.0% | 0.0% | 0.0% | 0.0% | 0.0% | 0.0% | 0.0% |
| Dumped pairs | 0.0% | 0.0% | 0.0% | 0.0% | 0.0% | 0.0% | 0.0% |

**C. Duplicates and contact ranges**

| | Proportion of read pairs | | | | | | |
|---|---|---|---|---|---|---|---|
| Number of input read pairs | 500K | 1M | 5M | 10M | 50M | 100M | 200M |
| Valid interaction | 58.0% | 58.0% | 57.9% | 57.9% | 57.9% | 58.0% | 57.9% |
| Valid interaction (remove duplicates) | 58.0% | 58.0% | 57.8% | 57.7% | 56.7% | 55.6% | 53.4% |
| Trans interaction | 35.8% | 35.7% | 35.6% | 35.5% | 34.9% | 34.2% | 32.9% |
| Cis interaction | 22.2% | 22.3% | 22.2% | 22.2% | 21.8% | 21.4% | 20.5% |
| Cis short-range interaction (<20kb) | 7.6% | 7.6% | 7.6% | 7.5% | 7.4% | 7.3% | 7.0% |
| Cis long-range interaction (>20kb) | 14.7% | 14.7% | 14.7% | 14.6% | 14.4% | 14.1% | 13.5% |

Note that the *cis* /*trans* judgement based on fragmentary sequences might be unreliable because insufficient continuity increases the trans fraction and instead decreases the cis fraction.

**Supplementary Table S7:** Quality control of the human GM12878 Hi-C libraries

| **Library preparation condition** | | |
|---|---|---|
| Cell fixation duration (min) | 10 | 30 |
| Amount of tissue for Hi-C reaction (cell number, ug DNA, or mg tissue) | $1.5 \times 10^6$ cells | |
| Restriction enzyme | HindIII | |
| Amount of Hi-C DNA used for library preparation (ug) | 2 | |
| PCR cycles | 6 | |
| Library size (bp) | 450 | 478 |
| Library yield (ng) | 140.4 | 112.2 |

| **Read pair category** | Proportion of read pairs | |
|---|---|---|
| Unique paired alignments | 72.2% | 76.0% |
| Unmapped pairs | 2.6% | 2.8% |
| Low quality pairs | 0.0% | 0.0% |
| Multiple pairs alignments | 15.6% | 14.4% |
| Pairs with singleton | 9.6% | 6.8% |
| Low quality singleton | 0.0% | 0.0% |
| Unique singleton alignments | 0.0% | 0.0% |
| Multiple singleton alignments | 0.0% | 0.0% |
| Reported pairs | 72.2% | 76.0% |

| **Hi-C read pair category** | Proportion of read pairs | |
|---|---|---|
| Valid interaction pairs | 66.4% | 59.4% |
| Dangling end pairs | 2.5% | 7.9% |
| Religation pairs | 2.7% | 7.8% |
| Self circle pairs | 0.5% | 0.9% |
| Single-end pairs | 0.0% | 0.0% |
| Filtered pairs | 0.0% | 0.0% |
| Dumped pairs | 0.0% | 0.0% |

### iconHi-C Protocol (ver. 1.0)



#### Step 1. (DAY 0) Preparation of cells/tissue

<table>
<tr><td rowspan="27">Cell/tissue preparation</td><td colspan="2"><u>Cultured cells and nucleated red blood cells</u></td></tr>
<tr><td>1</td><td>Collect $1 \times 10^7$ cells in a microtube (1.5 or 2.0 ml).</td></tr>
<tr><td>2</td><td>Centrifuge the cells (500 ×g, 5 min, 4°C), remove the supernatant, put on ice, and proceed immediately to fixation (Step 2.1a).</td></tr>
<tr><td colspan="2"><u>Tissues</u></td></tr>
<tr><td>1</td><td>Dissect and collect tissue in a microtube (1.5 or 2.0 ml) and freeze immediately in liquid nitrogen.<br>Note: Store the frozen tissue in an ultra-low temperature freezer (e.g., -80°C).</td></tr>
<tr><td>2</td><td>Pre-cool a mortar and a pestle in an ultra-low temperature freezer for at least 1 hr.</td></tr>
<tr><td>3</td><td>Pour liquid nitrogen into the pre-cooled mortar with a pestle.</td></tr>
<tr><td>4</td><td>Transfer tissue (up to 1 $cm^3$ in size) into the mortar and grind until it becomes a fine powder.</td></tr>
<tr><td>5</td><td>Transfer the tissue suspension, in liquid nitrogen, into a 50 ml tube pre-cooled with liquid nitrogen.</td></tr>
<tr><td>6</td><td>Close the screw cap loosely, put the tube in an ultra-low temperature freezer, and wait until the liquid nitrogen evaporates completely.</td></tr>
<tr><td>7</td><td>Proceed immediately to cell fixation (Step 2.1b) or store the powderized tissue in an ultra-low temperature freezer (e.g., -80°C) until use.</td></tr>
<tr><td colspan="2"><u>Alternative preparation method for soft tissues (e.g., liver, brain, testis, and embryonic tissues) using a frost-mill</u></td></tr>
<tr><td>1</td><td>Dissect and collect tissue in a microtube (1.5 or 2.0 ml) and freeze immediately in liquid nitrogen.<br>Note: Store the frozen tissue in an ultra-low temperature freezer (e.g., -80°C).</td></tr>
<tr><td>2</td><td>Pre-cool the Tokken stainless-steel tubes and bullets in liquid nitrogen.</td></tr>
<tr><td>3</td><td>Transfer tissue (up to 4 $mm^3$ in size) into each stainless-steel tube.<br>Notes: Break a large tissue into small pieces if necessary; Do not let the tissue thaw during the powderization process.</td></tr>
<tr><td>4</td><td>Put a bullet in the stainless-steel tube and close the screw cap.</td></tr>
<tr><td>5</td><td>Assemble the Tokken tube holder with three stainless-steel tubes.<br>Note: The tube holder should always be assembled with three stainless-steel tubes to balance the weight.</td></tr>
<tr><td>6</td><td>Submerge the tube holder (with tubes) in liquid nitrogen and wait ~1 min.</td></tr>
<tr><td>7</td><td>Take out the tube holder (with tubes) from liquid nitrogen, immediately place inside the Tokken acryl-grinder and vigorously shake the grinder for about 100 strokes to powderize the tissue.</td></tr>
<tr><td>8</td><td>Remove the bullet and collect the powderized tissue in a microtube (1.5 or 2.0 ml) pre-cooled in liquid nitrogen.<br>Note: Hand warm the lid of the deep-frozen microtube before closing its lid; otherwise, the tube can break easily.</td></tr>
<tr><td>9</td><td>Proceed immediately to cell fixation (Step 2.1a) or store the powderized tissue in an ultra-low temperature freezer (e.g., -80°C) until use.</td></tr>
</table>

#### Step 2. (DAY 0) Fixation of cells/tissue

Follow workflow a (for cultured cells, nucleated red blood cells, and tissue powderized using the Tokken frost-mill) or b (for tissue powderized using a mortar and a pestle).

<table>
<tr><td rowspan="8">Workflow a</td><td>1a</td><td>Prepare the fixing solution and put on ice.<br><table><tr><td><u>Fixing solution (10 ml)</u></td><td></td><td>(final)</td></tr><tr><td>16% formaldehyde solution</td><td>1 ml</td><td>(1%)</td></tr><tr><td>PBS (-)</td><td>15 ml</td><td></td></tr><tr><td>total</td><td>16 ml</td><td></td></tr></table>Note: Make fresh every time.</td></tr>
<tr><td>2a</td><td>Take a sample tube (microtube) out from the ice bucket or the ultra-low temperature freezer, immediately add 1-1.5 ml of the ice-cold fixing solution, and vortex mix until the cells/tissue powder is fully resuspended in the fixing solution.</td></tr>
<tr><td>3a</td><td>Incubate the cell-suspension for 10 min, or the tissue-suspension for 15-20 min in a heat block or a water bath set at 25°C.</td></tr>
<tr><td>4a</td><td>Quick spin the sample tube, add 1/20 volume of a 2.5 M glycine solution to quench the formaldehyde, vortex mix, and put on ice for ~ 1 min.</td></tr>
<tr><td>5a</td><td>Centrifuge the sample tube (1,000 ×g, 5 min, 4°C), remove the supernatant, add 1 ml of ice-cold PBS (-) and vortex mix.</td></tr>
<tr><td>6a</td><td>Repeat Step 2.5a.</td></tr>
<tr><td>7a</td><td>Resuspend the pellet in 1-1.5 ml of ice-cold PBS and 1) aliquot into microtubes (1.5 ml) at 1-2 $\times 10^6$ cells/tube, or, 2) aliquot the tissue suspension evenly into five to ten microtubes (1.5 ml).<br>Note: Aliquot a 4 $mm^3$-size tissue into five to ten microtubes; Adjust the number of microtubes proportionally for tissue at a different size.</td></tr>
<tr><td>8a</td><td>Centrifuge the sample tubes (1,000 ×g, 5 min, 4°C), remove the supernatant, and store in an ultra-low temperature freezer (e.g., -80°C) until use.</td></tr>
</table>

| | | |
|---|---|---|
| Workflow b | 1b | Prepare the fixing solution and put on ice.<br>Fixing solution (10 ml) — (final)<br>16% formaldehyde solution — 1 ml — (1%)<br>PBS (-) — 15 ml<br>total — 16 ml |
| | 2b | Take a sample tube (50 ml tube) out from the ultra-low temperature freezer, immediately add 10-15 ml of the ice-cold fixing solution, and vortex mix until the tissue powder is fully resuspended in the fixing solution. |
| | 3b | Incubate the tissue suspension for 15-20 min in a heat block or a water bath set at 25°C. |
| | 4b | Quick spin the sample tube, add 1/20 volume of a 2.5 M glycine solution to quench the formaldehyde, vortex mix, and put on ice for ~ 1 min. |
| | 5b | Centrifuge the sample tube (1,000 ×g, 5 min, 4°C), remove the supernatant, add 10 ml of ice-cold PBS (-), and vortex mix. |
| | 6b | Repeat Step 2.5b. |
| | 7b | Resuspend the pellet in 1-5 ml of ice-cold PBS and aliquot evenly into five to ten microtubes (1.5 ml).<br>Note: Aliquot a 4 $mm^3$-size tissue into five to ten microtubes; Adjust the number of microtubes proportionally for tissue at a different size. |
| | 8b | Centrifuge the sample tubes (1,000 ×g, 5 min, 4°C), remove the supernatant, and store in an ultra-low temperature freezer (e.g., -80°C) until use. |

##### Step 3. (DAY 0) Pre-determination of the amount of tissue to use for Hi-C

The pre-determination step quantitates the amount of DNA contained in a cell/tissue pellet prepared in Step 2.8a or 2.8b. Pre-determination should always be performed for tissue samples, but it is optional when the DNA content in a pellet is estimable,. e.g., for cultured cells and nucleated red blood cells.

| | | |
|---|---|---|
| Cell/tissue lysis | 1 | Take a microtube out from the ultra-low temperature freezer (from Step 2.8a or 2.8b), and immediately add 300 μl of lysis buffer (10 mM Tris-HCl pH8.0, 300 mM NaCl, 5 mM EDTA, 1% SDS). |
| | 2 | Add 10 μl of Proteinase K (20 mg/ml) to the sample tube, pipet mix, and incubate for ~16 hrs at 65°C, in a heat block, a water bath, or a hybridization oven. |
| | 3 | Add 5 μl of RNase A (10 mg/ml) to the sample tube, mix gently, and incubate in a thermal mixer (e.g., Eppendorf ThermoMixer C) for 20 min at 37°C, 950 rpm. |
| | 4 | Add 5 μl of Proteinase K (20 mg/ml) to the sample tube, mix gently, and incubate in a thermal mixer for 40 min at 55°C, 950 rpm. |
| DNA extraction and quantification | 5 | Take the sample tube out from the thermal mixer, let it cool down to room temperature, and proceed with Phenol-Chloroform DNA extraction. |
| | 6 | Add 300 μl of Phenol/Chloroform/Isoamyl alcohol solution and mix gently. |
| | 7 | Centrifuge the sample tube (16,000 ×g, 5 min, RT), and transfer ~270 μl of the aqueous phase into a new 1.5 ml microtube. |
| | 8 | Add 300 μl of TE/NaCl solution (10 mM Tris-HCl pH 8.0, 250 mM NaCl, 1 mM EDTA) to the sample tube containing Phenol, mix gently, and centrifuge again (16,000 ×g, 5 min, RT). |
| | 9 | Transfer ~300 μl of the aqueous phase into the microtube containing the first aqueous phase.<br>Note: The total volume of the aqueous phase will be ~570 μl. |
| | 10 | Add 1 μl of glycogen solution (20 mg/ml) to the collected aqueous phase and mix gently. |
| | 11 | Add 600 μl of 2-Propanol to the collected aqueous phase and gently mix until the solution becomes homogeneous. |
| | 12 | Centrifuge the sample tube (20,000 ×g, 30 min, 4°C). |
| | 13 | Decant the supernatant, add 1 ml of 70% EtOH, and mix gently to rinse the DNA pellet. |
| | 14 | Centrifuge the sample tube (20,000 ×g, 10 min, 4°C). |
| | 15 | Decant the supernatant and centrifuge the sample tube again (20,000 ×g, 5 min, 4°C). |
| | 16 | Remove the supernatant completely with a P-100 or P-200 pipet, and keep the lid of the sample tube open for ~1 min to allow the ethanol to evaporate.<br>Note: Do not over dry the pellet. |
| | 17 | Dissolve DNA in 30-50 μl of TE. |
| | 18 | Quantitate the DNA using 1 μl of the DNA sample with Qubit dsDNA High Sensitivity Kit, and calculate the total amount of DNA extracted from a cell/tissue pellet (from Step 2.8a or 2.8b). |
| | 19 | Determine the amount of cell/tissue to use for Hi-C.<br>Note: Use a cell/tissue pellet that contains 2-10 μg of DNA. |

##### Step 4. (DAY 1) Restriction enzyme digestion

The number of cells to use for Hi-C is determined based on the amount of DNA. Use $1\times10^6$ cells for an animal with the genome size of 3-3.5 Gb (e.g., human and mouse), $2\times10^6$ cells for an animal with the genome size of 1-1.5 Gb-size (e.g., chicken and western clawed frog), or a cell/tissue that contains 2-10 µg of DNA.
Follow cells/tissue resuspension steps as written, i.e., vortex mix or pipet mix.

<table>
<tr><td rowspan="18">Cell/tissue permeabilization</td><td>1</td><td>Prepare the permeabilization buffer 1 (PB1).
<table>
<tr><th>Permeabilization buffer 1</th><th></th><th>(final)</th></tr>
<tr><td>1 M Tris-HCl (pH 8.0)</td><td>400 µl</td><td>(10 mM)</td></tr>
<tr><td>5 M NaCl</td><td>80 µl</td><td>(10 mM)</td></tr>
<tr><td>10% (w/v) NP-40</td><td>800 µl</td><td>(0.2%)</td></tr>
<tr><td>H2O</td><td>38.72 ml</td><td></td></tr>
<tr><td>Total</td><td>40 ml</td><td></td></tr>
</table>
Note: Filtrate and store at 4°C.</td></tr>
<tr><td>2</td><td>Take (400 µl ×n) + 5% extra volume of PB1 in a new tube, add 1/100 vol of a proteinase inhibitor cocktail (PI), and put on ice.<br>Note: Add PI to the PB1 just before use.</td></tr>
<tr><td>3</td><td>Take the frozen cells/tissue out from the freezer (from Step 2.8a or 2.8b), immediately add 400 µl of PB1 (with PI), vortex mix, and follow Step 4.4a (for cultured cells, nucleated red blood cells, and tissue powderized using a mortar and a pestle) or Step 4.4b (for tissue powderized using the Tokken frost-mill).</td></tr>
<tr><td>4a</td><td>Incubate the sample tube on ice for 20 min with periodical mixing every 5-10 min.</td></tr>
<tr><td>4b</td><td>Homogenize the tissue by ~20 strokes of homogenization in a douncer with the tight pestle B on ice, and transfer the homogenate into a 1.5 ml microtube.</td></tr>
<tr><td>5</td><td>Centrifuge the sample tube (2,000 ×g, 3-5 min, 4°C).</td></tr>
<tr><td>6</td><td>Remove 300 µl of the supernatant using a P-1000 pipet, centrifuge the sample tube again (2,000 ×g, 3-5 min, 4°C), and remove the remaining supernatant using a P-100 or a P-200 pipet.</td></tr>
<tr><td>7</td><td>Prepare the permeabilization buffer 2 (PB2) at room temperature.
<table>
<tr><th>Permeabilization buffer 2</th><th></th><th>(× )</th><th>(final)</th></tr>
<tr><td>10X NEBuffer 2</td><td>25 µl</td><td>µl</td><td>(1X)</td></tr>
<tr><td>20 mg/ml BSA</td><td>1.25 µl</td><td>µl</td><td>(0.1 mg/ml)</td></tr>
<tr><td>10% (w/v) SDS</td><td>7.5 µl</td><td>µl</td><td>(0.3%)</td></tr>
<tr><td>PI (100X)</td><td>2.5 µl</td><td>µl</td><td>(1X)</td></tr>
<tr><td>H2O</td><td>213.75 µl</td><td>µl</td><td></td></tr>
<tr><td>Total</td><td>250 µl</td><td>µl</td><td></td></tr>
</table>
Notes: No need to add BSA if the buffer already contains BSA, e.g., NEBuffer 2.1; Make fresh every time; Make 5% extra volume.</td></tr>
<tr><td>8</td><td>Add 250 µl of PB2 to the sample tube and pipet mix.</td></tr>
<tr><td>9</td><td>Incubate/shake the sample tube in a thermal mixer for 10 min at 37°C, 950 rpm.</td></tr>
<tr><td>10</td><td>Add 28 µl of 20% TritonX-100 to the sample tube.</td></tr>
<tr><td>11</td><td>Incubate/shake the sample tube in a thermal mixer for 10 min at 37°C, 950 rpm.</td></tr>
<tr><td>12</td><td>Take 8 µl (3%) aliquot in a new 1.5 ml microtube as a pre-digest DNA control (ctr-1), add 42 µl of DNA-preparation buffer and store at -20°C until the end of DAY2.
<table>
<tr><th>DNA-preparation buffer</th><th></th><th>(final)</th></tr>
<tr><td>5 M NaCl</td><td>250 µl</td><td>(250 mM)</td></tr>
<tr><td>0.5 M EDTA</td><td>50 µl</td><td>(5 mM)</td></tr>
<tr><td>10% (w/v) SDS</td><td>500 µl</td><td>(1%)</td></tr>
<tr><td>H2O</td><td>4.2 ml</td><td></td></tr>
<tr><td>total</td><td>5 ml</td><td></td></tr>
</table>
Notes: Store DNA-preparation buffer at room temperature; Warm the buffer at 37°C in case the SDS precipitates during storage.</td></tr>
<tr><td>13</td><td>Centrifuge the sample tube (2,000 ×g, 3-5 min, 4°C).</td></tr>
<tr><td>14</td><td>Remove 200 µl of the supernatant using a P-200 pipet, centrifuge the sample tube again (2,000 ×g, 3-5 min, 4°C), and remove the remaining supernatant using a P-100 or a P-200 pipet.</td></tr>
<tr><td>15</td><td>Prepare the washing buffer (WB) with NEBuffer DpnII (for DpnII digestion) or NEBuffer 2 (for HindIII digestion).
<table>
<tr><th>Washing Buffer</th><th></th><th>(× )</th><th>(final)</th></tr>
<tr><td>10X NEBuffer</td><td>50 µl</td><td>µl</td><td>(1X)</td></tr>
<tr><td>20 mg/ml BSA</td><td>1.25 µl</td><td>µl</td><td>(0.1 mg/ml)</td></tr>
<tr><td>20% (w/v) TritonX-100</td><td>1.25 µl</td><td>µl</td><td>(0.05%)</td></tr>
<tr><td>H2O</td><td>447.5 µl</td><td>µl</td><td></td></tr>
<tr><td>total</td><td>500 µl</td><td>µl</td><td></td></tr>
</table>
Notes: No need to add BSA if the buffer already contains BSA, e.g., NEBuffer 2.1; Do not use NEBuffer 3.1 for DpnII because non-specific cleavage (star activity) may be induced; Make 5% extra volume.</td></tr>
<tr><td>16</td><td>Add 500 µl of WB to the sample tube and vortex mix.</td></tr>
<tr><td>17</td><td>Centrifuge the sample tube (2,000 ×g, 3-5 min, 4°C).</td></tr>
</table>

<table>
<tr><td></td><td>18</td><td>Remove 400 µl of the supernatant using a P-1000 pipet, centrifuge the sample tube again (2,000 ×g, 3-5 min, 4°C), and remove the remaining supernatant using a P-100 pipet or a P-200 pipet.</td></tr>
<tr><td rowspan="3">Restriction enzyme digestion</td><td>19</td><td>Prepare the restriction-enzyme mix on ice.<br><table>
<tr><th>Restriction-enzyme mix</th><th></th><th>(× )</th><th>(final)</th></tr>
<tr><td>10X NEBuffer</td><td>20 µl</td><td>µl</td><td>(1X)</td></tr>
<tr><td>20 mg/ml BSA</td><td>1 µl</td><td>µl</td><td>(0.1 mg/ml)</td></tr>
<tr><td>RE (50 U/ul)</td><td>8 µl</td><td>µl</td><td>(400 U)</td></tr>
<tr><td>20% Triton X-100</td><td>0.5 µl</td><td>µl</td><td>(0.05%)</td></tr>
<tr><td>H2O</td><td>170.5 µl</td><td>µl</td><td></td></tr>
<tr><td>total</td><td>200 µl</td><td>µl</td><td></td></tr>
</table><br>Notes: No need to add BSA if the buffer already contains BSA, e.g., NEBuffer 2.1; Do not use NEBuffer 3.1 for DpnII because non-specific cleavage (star activity) may be induced; For restriction enzymes at different concentration, e.g., HindIII at 100U/µl, adjust the volume with H2O; The restriction-enzyme mix may or may not contain Triton X-100; Make 5% extra volume.</td></tr>
<tr><td>20</td><td>Add 200 µl of restriction-enzyme mix to the sample tube and pipet mix.</td></tr>
<tr><td>21</td><td>Incubate the sample tube in a thermal mixer for ~16 hrs at 37°C, 1,100 rpm.</td></tr>
</table>

#### Step 5. (DAY 2) DNA fill-in and ligation

Follow cells/tissue resuspension steps as written, i.e., vortex mix or pipet mix.

**DNA fill-in reaction**

1 Take 6 µl (3%) aliquot in a new 1.5 ml microtube as a digested-DNA control (ctr-2), add 44 µl of DNA-preparation buffer and store at -20°C until the end of DAY 2.

2 Centrifuge the sample tube (2,000 ×g, 3-5 min, 4°C).

3 Remove 100 µl of the supernatant using a P-200 pipet, centrifuge the sample tube again (2000 ×g, 3-5 min, 4°C), and remove the remaining supernatant using a P-200 pipet or a P-100 pipet.

4 Prepare the washing buffer (WB).

| Washing buffer | | (× | ) | (final) |
|---|---|---|---|---|
| 10X NEBuffer2 | 50 µl | | µl | (1X) |
| 20 mg/ml BSA | 1.25 µl | | µl | (0.1 mg/ml) |
| 20% (w/v) TritonX-100 | 1.25 µl | | µl | (0.05%) |
| H2O | 447.5 µl | | µl | |
| total | 500 µl | | µl | |

Notes: No need to add BSA if the buffer already contains BSA, e.g., NEBuffer 2.1; Prepare enough buffer to perform four wash cycles or two wash cycles (at Steps 5.5 and 5.13), for the DpnII-digested sample or the HindIII-digested sample respectively.

5 Add 500 µl of WB to the sample tube and vortex mix.

6 Centrifuge the sample tube (2,000 ×g, 3-5 min, 4°C).

7 Remove 400 µl of the supernatant using a P-1000 pipet, centrifuge the sample tube again (2,000 ×g, 3-5 min, 4°C), and remove the remaining supernatant using a P-100 pipet or a P-200 pipet.

Note: Repeat Steps 5.5-5.7 twice more for the DpnII sample (total of three wash cycles).

8 Prepare the DNA fill-in mix on ice.

| DNA fill-in mix for the DpnII-digested sample | | (× | ) | (final) |
|---|---|---|---|---|
| 10X NEBuffer 2 | 10 µl | | µl | (1X) |
| 1 mM dCTP | 1.5 µl | | µl | (15 µM) |
| 1 mM dGTP | 1.5 µl | | µl | (15 µM) |
| 1 mM dTTP | 1.5 µl | | µl | (15 µM) |
| 0.4 mM biotin-14-dATP | 3.75 µl | | µl | (15 µM) |
| Klenow DNA polymerase (5 U/µl) | 6 µl | | µl | (30 U) |
| 20% TritonX-100 | 0.25 µl | | µl | (0.05%) |
| H2O | 75.5 µl | | µl | |
| Total | 100 µl | | µl | |

| DNA fill-in mix for the HindIII-digested sample | | (× | ) | (final) |
|---|---|---|---|---|
| 10X NEBuffer 2 | 10 µl | | µl | (1X) |
| 1 mM dATP | 1.5 µl | | µl | (15 µM) |
| 1 mM dGTP | 1.5 µl | | µl | (15 µM) |
| 1 mM dTTP | 1.5 µl | | µl | (15 µM) |
| 0.4 mM biotin-14-dCTP | 3.75 µl | | µl | (15 µM) |
| Klenow DNA polymerase (5 U/µl) | 3 µl | | µl | (15 U) |
| 20% TritonX-100 | 0.25 µl | | µl | (0.05%) |
| H2O | 78.5 µl | | µl | |
| Total | 100 µl | | µl | |

Note: The DNA fill-in mix may or may not contain Triton X-100.

9 Add 100 µl of DNA fill-in mix to the sample tube and pipet mix.

10 Incubate the sample tube in a thermal mixer for 20 min at 25°C, 1,100 rpm.

11 Centrifuge the sample tube (1500 ×g, 3-5 min, 4°C).

12 Remove the supernatant using a P-100 pipet or a P-200 pipet.

13 Add 500 µl of WB to the sample tube and vortex mix.

14 Centrifuge the sample tube (2,000 ×g, 3-5 min, 4°C).

15 Remove 400 µl of the supernatant using a P-1000 pipet, centrifuge the sample tube again (2,000 ×g, 3-5 min, 4°C), and remove the remaining supernatant using a P-100 pipet or a P-200 pipet.

| | | |
|---|---|---|
| Ligation reaction | 16 | Prepare the ligation mix on ice. |
| | 17 | Add 100 μl of ligation mix to the sample tube and pipet mix. |
| | 18 | Incubate the sample tube in a thermal mixer for 4-6 hrs at 16°C, 1,100 rpm. |

| Ligation mix for the DpnII-digested sample | | (× ) | (final) |
|---|---|---|---|
| T4 DNA ligase buffer (10X) | 10 μl | μl | (1X) |
| T4 DNA ligase (2,000 U/ul) | 2 μl | μl | (4,000 CEU) |
| 20% TritonX-100 | 0.25 μl | μl | (0.05%) |
| H2O | 87.75 μl | μl | |
| total | 100 μl | μl | |

Notes: Prepare T4 DNA ligase buffer in small aliquots and store them at -20°C when the buffer is thawed for the first time; Always use an aliquot that is not frozen and thawed repeatedly; The ligation mix may or may not contain Triton X-100; Use 2,000 CEU of T4 DNA ligase for the HindIII-digested sample.

**Step 6.1. (DAY 2) DNA purification**

| | | |
|---|---|---|
| DNA purification | 1 | Prepare the DNA-extraction mix. |
| | 2 | Add 200 μl of DNA−extraction mix to the sample tube (from Step 5.18) and mix gently. Add 200 μl of DNA-extraction mix and 50 μl of H2O to the control tubes also (ctr-1 from Step 4.12 and ctr-2 from Step 5.1). Note: The total amount of the mixture will be 300 ul. |
| | 3 | Incubate the sample tubes in a thermal mixer for ~16 hrs at 65°C, 350 rpm. |

| DNA-extraction mix | | (× ) | (final) |
|---|---|---|---|
| 1M Tris-HCl (pH 8.0) | 2 μl | μl | (10 mM) |
| 0.5M EDTA | 2 μl | μl | (5 mM) |
| 10% SDS | 30 μl | μl | (1.5%) |
| 5M NaCl | 15 μl | μl | (375 mM) |
| 20 mg/ml Proteinase K | 10 μl | μl | (1 mg/ml) |
| H2O | 141 μl | μl | |
| total | 200 μl | μl | |

Note: Prepare DNA-extraction mix at room temperature.

##### Step 6.2. (DAY 3) DNA purification

| | | |
|---|---|---|
| DNA purification | 1 | Take the sample tubes out from the thermal mixer (from Step 6.1.3) and let them cool down to room temperature. |
| | 2 | Add 5 µl of RNase A (10 mg/ml) to the sample tube, mix gently, and incubate in a thermal mixer for 20 min at 37°C, 800 rpm. |
| | 3 | Add 5 µl of Proteinase K (20 mg/ml) to the sample tube, mix gently, and incubate in a thermal mixer for 2 hrs at 55°C, 800 rpm. |
| | 4 | Take out the sample tubes from the thermal mixer, let them cool down to room temperature, and proceed with DNA extraction. |
| | 5 | Add 300 µl of Phenol/Chloroform/Isoamyl alcohol solution to the sample tube and mix gently. |
| | 6 | Centrifuge the sample tube (16,000 ×g, 5 min, RT), and transfer ~270 µl of the aqueous phase into a new 1.5 ml microtube. |
| | 7 | Add 300 µl of TE/NaCl solution (10 mM Tris-HCl pH 8.0, 250 mM NaCl, 1 mM EDTA) to the sample tube containing Phenol, mix gently, and centrifuge again (16,000 ×g, 5 min, RT). |
| | 8 | Transfer ~300 µl of the aqueous phase into the microtube containing the first aqueous phase.<br>Note: The total volume of the collected aqueous phase will be ~570 µl. |
| | 9 | Add 1 µl of glycogen solution (20 mg/ml) to the collected aqueous phase and mix gently. |
| | 10 | Add 600 µl of 2-propanol to the collected aqueous phase and mix gently until the solution becomes homogeneous. |
| | 11 | Centrifuge the sample tube (20,000 ×g, 30 min, 4°C). |
| | 12 | Decant the supernatant, add 1 ml of 70% EtOH, and mix gently to rinse the DNA pellet. |
| | 13 | Centrifuge the sample tube (20,000 ×g, 10 min, 4°C). |
| | 14 | Decant the supernatant and centrifuge the sample tube again (20,000 ×g, 5 min, 4°C). |
| | 15 | Remove the supernatant completely with a P-100 or P-200 pipet, and keep the lid open for ~1 min to allow the ethanol to evaporate.<br>Note: Do not over dry the pellet. |
| | 16 | Dissolve the Hi-C DNA in 30 µl of EB and the control DNAs (ctr-1 and ctr-2) in 10 µl of EB.<br>Notes: Mix gently to avoid shearing of the DNA; Store DNA samples at 4°C or -20°C. |
| | 17 | Quantitate the DNA using 1 µl of the DNA sample with the Qubit dsDNA High Sensitivity Kit. |

##### Step 7. (DAY 3) Hi-C DNA QC (QC1)

| | | |
|---|---|---|
| Hi-C DNA QC (QC1) | 1 | Take 1-2 µl of the DNA sample (ctr-1, ctr-2, and Hi-C DNA) from Step 6.2.16 in a new PCR tube and adjust the concentration to 2-20 ng/µl with EB.<br>Note: The concentration of the DNA samples in a trio (ctr-1, ctr-2, and Hi-C DNA) should be within two-fold of difference. |
| | 2 | Analyze 1 µl of the DpnII-digested DNA samples (ctr-2 and Hi-C DNA) using the Agilent Bioanalyzer with the DNA High Sensitivity chip, or analyze 1 µl of the HindIII-digested DNA samples (ctr-1, ctr-2, and Hi-C DNA) using the Agilent TapeStation with the genomic tape.<br>Note: Ctr-1 (pre-digested DNA) is >50 kb in size and cannot be analyzed using the Agilent Bioanalyzer with the DNA High Sensitivity chip. |
| | 3 | Check the pattern of size shift between samples and the level of DNA degradation.<br>Note: Only the qualified Hi-C DNA, showing the expected pattern of size shift with no or minimum degree of DNA degradation, is used for the preparation of the Hi-C library. |
| | 4 | Store Hi-C DNA at -20°C or proceed to Step 8. |

##### Step 8. (DAY 3) Removal of biotin from un-ligated DNA ends

| | | |
|---|---|---|
| Removal of biotin from un-ligated DNA ends | 1 | Take 250 ng-2 µg of Hi-C DNA (from Step 7.4) in a 0.2 ml PCR tube and adjust the total volume to 30 µl with H2O. |
| | 2 | Prepare the T4-DNA-polymerase mix on ice. |
| | 3 | Add 20 µl T4-DNA-polymerase mix to the Hi-C DNA, mix gently and incubate in a PCR machine for 30 min at 37°C, and 15 min at 75°C. |

| T4-DNA-polymerase mix for the DpnII-digested sample | | (× ) | (final) |
|---|---|---|---|
| 10X NEBuffer 2 | 5 µl | µl | (1X) |
| 20 mg/ml BSA | 0.5 µl | µl | (0.2 mg/ml) |
| 10 mM GTP | 0.5 µl | µl | (100 µM) |
| T4 DNA pol (3 U/ul) | 1.67 µl | µl | (5 U) |
| H2O | 12.33 µl | µl | |
| Total | 20 µl | µl | |

| T4-DNA-polymerase mix for the HindIII-digested sample | | (× ) | (final) |
|---|---|---|---|
| 10X NEBuffer 2 | 5 µl | µl | (1X) |
| 20 mg/ml BSA | 0.5 µl | µl | (0.2 mg/ml) |
| 10 mM ATP | 0.5 µl | µl | (100 µM) |
| 10 mM GTP | 0.5 µl | µl | (100 µM) |
| T4 DNA pol (3 U/ul) | 0.84 µl | µl | (2.5 U) |
| H2O | 12.66 µl | µl | |
| Total | 20 µl | µl | |

##### Step 9. (DAY 3) Fragmentation and size selection of the Hi-C DNA

| | | |
|---|---|---|
| Fragmentation and size selection of the Hi-C DNA | 1 | Turn on the Covaris (S220 or E220). |
| | 2 | Transfer the entire reaction from Step 8.3 into a Covaris microTUBE and add 80 µl of TE buffer.<br>Note: The total volume will be 130 µl. |
| | 3 | Perform sonication (Duty factor: 5%, Peak incident power: 175, Cycles per burst: 200, Time: 60 sec ×2, temperature: 7°C). |
| | 4 | Transfer 120 µl of the fragmented DNA into a 1.5 ml microtube. |
| | 5 | Add 72 µl (×0.6 amount) of AMPure XP beads to the fragmented DNA, vortex and incubate 5 min at room temperature.<br>Note: Bead-bound DNA larger than 600 bp is removed at Steps 9.5-9.7. |
| | 6 | Quick spin the sample tube, put on the magnetic and wait until the supernatant becomes clear. |
| | 7 | Transfer the supernatant into a new 1.5 ml microtube. |
| | 8 | Add 108 µl (×0.9 amount) of AMPure XP beads to the supernatant, vortex mix, and wait 5 min at room temperature.<br>Note: Bead-bound DNA larger than 150 bp is collected at Steps 9.8-9.9. |
| | 9 | Quick spin the sample tube, put on the magnetic and wait until the supernatant becomes clear. |
| | 10 | Remove the supernatant using a P-200 pipet while the sample tube is still on the magnet. |
| | 11 | Add 200 µl of 80% EtOH to the beads while the sample tube is still on the magnet and wait 30 sec. |
| | 12 | Remove EtOH with P-200 pipet while the sample tube is still on the magnet. |
| | 13 | Repeat the washing cycle (Steps 9.9-9.10). |
| | 14 | Quick spin the sample tube, put on the magnet and remove the residual EtOH completely using a P-10 or a P-10 pipet. |
| | 15 | Air dry the beads for ~1 min at room temperature with the lid kept open. |
| | 16 | Add 60 µl EB to the beads, vortex mix and incubate 2 min at room temperature. |
| | 17 | Quick spin the sample tube, put on the magnet and collect the eluate in a new PCR tube. |

##### Step 10. (DAY 3) Enrichment of biotin-containing DNA

| | | |
|---|---|---|
| Enrichment of the biotin-containing DNA | 1 | Prepare the 2X binding-and-washing buffer (BWB).<br>2X binding-and-washing buffer — (final)<br>1 M Tris-HCl (pH 7.5): 400 µl (10 mM)<br>0.5 M EDTA (pH 8.0): 80 µl (1 mM)<br>5 M NaCl: 16 ml (2 M)<br>10% Tween20: 80 µl (0.02%)<br>H2O: 23.44 ml<br>total: 40 ml<br>Note: Store at room temperature. |
| | 2 | Prepare 1X BWB by diluting the 2X BWB with H2O.<br>Note: Store at room temperature. |
| | 3 | Mix the bottle of the streptavidin beads and transfer (25 µl × n) + 5% extra volume of beads into a new 1.5 ml microtube. |
| | 4 | Put the tube on the magnetic and wait until the supernatant becomes clear. |
| | 5 | Remove the supernatant, take off the tube from the magnet, add 1 ml 1X BWB (prepared in Step 10.2), and vortex mix. |
| | 6 | Quick spin the tube, put on the magnetic and wait until the supernatant becomes clear. |
| | 7 | Remove the supernatant, take off the tube from the magnet, add (60 µl × n) + 5% extra volume of 2X BWB, and pipet mix. |
| | 8 | Add 60 µl beads (in 2X BWB) to the size-selected Hi-C DNA (prepared in Step 9.16) and vortex mix. |
| | 9 | Incubate the sample tube in a thermal mixer for 15 min at 20°C with periodical mixing for 10 sec at 2,000 rpm every 3 min. |
| | 10 | Quick spin the sample tube, put on the magnetic and wait until the supernatant becomes clear. |
| | 11 | Remove the supernatant, take off the sample tube from the magnet, add 100 µl of 1X BWB, and vortex mix. |
| | 12 | Quick spin the sample tube, put on the magnetic and wait until the supernatant becomes clear. |
| | 13 | Repeat the bead washing cycles (Steps 10.11-12) three more times (perform four times in total). |
| | 14 | Remove the supernatant, add 100 µl of EB, and vortex mix. |
| | 15 | Quick spin the sample tube, put on the magnetic and wait until the supernatant becomes clear. |
| | 16 | Remove the supernatant, add 100 µl of EB with the sample tube kept on the magnet. |
| | 17 | Remove the supernatant, take off the sample tube from the magnet, quick spin, and put the sample tube back on the magnet. |
| | 18 | Remove the residual EB with a P-10 or a P-20 pipet. |
| | 19 | Add 50 µl of EB and store the sample tube at 4°C until the next day. |

##### Step 11. (DAY 4) Hi-C library preparation

Library preparation is performed using the KAPA LTP DNA library kit but in a 1/5 reaction volume of the original protocol and with an additional step for PCR cycle pre-determination.

| Stage | Step | Procedure |
|---|---|---|
| End repair | 1 | Prepare the end-repair mix on ice |

| End-repair mix | | (× ) |
|---|---|---|
| 10X KAPA End Repair Buffer | 1.4 µl | µl |
| KAPA End Repair Enzyme Mix | 1.0 µl | µl |
| H2O | 11.6 µl | µl |
| Total volume | 14 µl | µl |

| Stage | Step | Procedure |
|---|---|---|
| End repair | 2 | Quick spin the tube (from Step 10.19), put on the magnetic and wait until the supernatant becomes clear. |
| End repair | 3 | Remove the supernatant, add 14 µl end-repair mix to the beads, pipet mix, and incubate for 30 min at 20°C (followed by a hold at 4°C) in a PCR machine. |
| End repair | 4 | Wash the beads as described in Steps 10.11-10.18 (4 times in 1X BWB, followed by 2 times in EB), close the lid, and put on ice. |
| A-tailing | 5 | Prepare the A-tailing mix on ice. |

| A-tailing mix | | (× ) |
|---|---|---|
| 10X KAPA A-Tailing Buffer | 1.0 µl | µl |
| KAPA A-Tailing Enzyme | 0.6 µl | µl |
| H2O | 8.4 µl | µl |
| Total volume | 10 µl | µl |

| Stage | Step | Procedure |
|---|---|---|
| A-tailing | 6 | Add 10 µl A-tailing mix to the beads (from Step 11.4), pipet mix, and incubate 30 min at 30°C (followed by a hold at 4°C) in a PCR machine. |
| A-tailing | 7 | Wash the beads as described in Steps 10.10-10.18 (4 times in 1X BWB, followed by 2 times in EB), close the lid, and put on ice. |
| Adapter ligation | 8 | Prepare the ligation-buffer mix (without the ligase enzyme) on ice |

| Ligation-buffer mix | | (× ) |
|---|---|---|
| 5X KAPA Ligation Buffer | 2.0 µl | µl |
| H2O | 6.0 µl | µl |
| Total volume | 8 µl | µl |

| Stage | Step | Procedure |
|---|---|---|
| Adapter ligation | 9 | Add 8 µl of ligation-buffer mix to the beads (from Step 11.7), add 1 µl of 1 µM Illumina TruSeq compatible adapter, and pipet mix. |
| Adapter ligation | 10 | Add 1 µl KAPA T4 DNA ligase, pipet mix, and incubate for 15 min at 20°C (followed by a hold at 4°C) in a PCR machine. |
| Adapter ligation | 11 | Wash the beads as described in Steps 10.10-10.18 (4 times in 1X BWB, followed by 2 times in EB), close the lid, and put on ice. |
| Pre-PCR (releasing DNA off the streptavidin beads) | 12 | Prepare the pre-PCR mix. |

| Pre-PCR mix | | (× ) |
|---|---|---|
| 2X KAPA HiFi Ready Mix | 10 µl | µl |
| 10 µM TPC mix | 0.9 µl | µl |
| H2O | 9.1 µl | µl |
| Total volume | 20 µl | µl |

| Stage | Step | Procedure |
|---|---|---|
| Pre-PCR (releasing DNA off the streptavidin beads) | 13 | Add 20 µl pre-PCR mix to the beads (from Step 11.11), pipet mix and perform 4 cycles of PCR amplification at, 98°C 45 sec, 4 cycles of (98°C 15 sec, 60°C 30 sec, 72°C 30 sec), 72°C 1 min, and a hold at 4°C. |
| | 14 | Put the sample tube on the magnetic and wait until the supernatant becomes clear. |
| | 15 | Transfer the supernatant to a new PCR tube, add 20 µl (×1 volume) of AMPure XP beads, vortex mix, and wait 5 min at room temperature. |
| | 16 | Quick spin the sample tube, put on the magnetic and wait until the supernatant becomes clear. |
| | 17 | Remove the supernatant using a P-200 pipet while the sample tube is still on the magnet. |
| | 18 | Add 200 µl of 80% EtOH to the beads while the sample tube is still on the magnet and wait 30 sec. |
| | 19 | Remove EtOH with a P-200 pipet while the sample tube is still on the magnet. |
| | 20 | Repeat the washing cycle (Steps 11.18-11.19). |
| | 21 | Quick spin the sample tube, put on the magnet and remove the residual EtOH completely using a P-10 or a P-20 pipet. |
| | 22 | Air dry the beads for ~1 min at room temperature with the lid kept open. |
| | 23 | Add 11 µl EB to the beads, vortex mix, and incubate 2 min at room temperature. |
| | 24 | Quick spin the sample tube, put on the magnet and collect the eluate in a new PCR tube. |
| PCR cycle pre-determination | 25 | Prepare the real-time PCR mix. |

| Real-time-PCR mix | | (× ) |
|---|---|---|
| 2X KAPA HiFi HS real-time Mix | 5 µl | µl |
| 10 µM TPC mix | 0.35 µl | µl |
| H2O | 3.15 µl | µl |
| Total volume | 8.5 µl | µl |

| Stage | Step | Procedure |
|---|---|---|
| PCR cycle pre-determination | 26 | Dispense 8.5 µl real-time-PCR mix in a well of a PCR plate (384 well or 96 well) and add 1.5 µl of the ligated DNA (from Step 11.24). |
| | 27 | Dispense 10 µl of each Fluorescence Standards in the same plate in separate wells. |

| Section | Step | Procedure |
|---|---|---|
| | 28 | Run real-time PCR in a ROX minus condition at, 98°C 45 sec, 20 cycles of (98°C 15 sec, 60°C 30 sec, 72°C 30 sec), 72°C 1 min. |
| | 29 | Determine the threshold-PCR-cycle (Ct) reaching Fluorescence Standard 1 but not exceeding Fluorescence Standard 2. Notes: DNA for the library QC (QC2) is prepared by amplifying a small aliquot (1 µl) of the pre-PCR product (from Step 11.24) with (Ct +3) cycles of PCR. Hi-C library for sequencing is prepared by amplifying the remaining 8.5 µl of the pre-PCR product (from Step 11.24) with the pre-determined Ct cycle. |
| Amplification of the library for Hi-C library QC (QC2) | 30 | Prepare the PCR-mix for Hi-C library QC (QC2). |

| QC2-PCR mix | | (× ) |
|---|---|---|
| 2X KAPA HiFi HotStart Ready Mix | 5 µl | µl |
| 10 µM TPC | 0.45 µl | µl |
| H2O | 3.55 µl | µl |
| Total volume | 9 µl | µl |

| Section | Step | Procedure |
|---|---|---|
| Amplification of the library for Hi-C library QC (QC2) | 31 | Dispense 9 µl of the 'QC2-PCR mix' in a PCR tube, add 1 µl of the pre-PCR product (from Step 11.24), and perform PCR amplification at, 98°C 45 sec, (Ct +3) cycles of (98°C 15 sec, 60°C 30 sec, 72°C 30 sec), 72°C 1 min, and a hold at 4°C. |
| | 32 | Add 10 µl (×1 volume) of AMPure XP beads, vortex mix, and wait 5 min at room temperature. |
| | 33 | Follow Steps 11.16-11.22 to purify the DNA. |
| | 34 | Add 5 µl EB to the beads, vortex mix, and incubate 2 min at room temperature. |
| | 35 | Quick spin the sample tube, put on the magnet and collect the eluate in a new PCR tube. |
| | 36 | Quantitate the DNA using 1 µl of the DNA sample with the Qubit dsDNA High Sensitivity Kit. |
| | 37 | Proceed to the Hi-C library QC (QC2) at Step 11. |
| Library amplification | 38 | Prepare a PCR mix for Hi-C library amplification. |

| Library-amplification-PCR mix | | (× ) |
|---|---|---|
| 2X KAPA HiFi HotStart Ready Mix | 10 µl | µl |
| 10 µM TPC | 0.9 µl | µl |
| H2O | 0.6 µl | µl |
| Total volume | 11.5 µl | µl |

| Section | Step | Procedure |
|---|---|---|
| Library amplification | 39 | Dispense 11.5 µl of the 'Library-amplification-PCR mix' to the tube containing the pre-PCR product (from Step 11.24), and perform PCR amplification at, 98°C 45 sec, (pre-determined Ct) cycles of (98°C 15 sec, 60°C 30 sec, 72°C 30 sec), 72°C 1 min, and a hold at 4°C. |
| | 40 | Add 20 µl (×1 volume) of AMPure XP beads, vortex mix, and wait 5 min at room temperature. |
| | 41 | Purify the DNA following Steps 11.16-11.22. |
| | 42 | Add 30 µl EB to the beads, vortex mix, and incubate 2 min at room temperature. |
| | 43 | Quick spin the sample tube, put on the magnet and collect the eluate in a new microtube (1.5 ml). |
| | 44 | Quantitate the DNA using 1 µl of the DNA sample with the Qubit dsDNA High Sensitivity Kit. |
| | 45 | Analyze the size-distribution using 1 µl of the library DNA with Agilent TapeStation High Sensitivity D1000 tape. |

#### Step 12. (DAY 4) Quality control of the Hi-C library (QC2)

| Section | Step | Procedure |
|---|---|---|
| Hi-C library QC (QC2) | 1 | Prepare the restriction-mix for QC2 with or without the restriction enzyme. |

| QC2-restriction mix | | (× ) |
|---|---|---|
| 10X buffer | 1 µl | µl |
| RE (10 U/ul) or H2O | 0.3 µl | µl |
| 20 mg/ml BSA | 0.05 µl | µl |
| H2O | µl | µl |
| Total volume | ul | µl |

Note: Cut the DpnII-digested library with ClaI, and the HindIII-digested library with NheI.

| Section | Step | Procedure |
|---|---|---|
| Hi-C library QC (QC2) | 2 | Dispense 'QC2-restriction mix' (with or without the RE) to a PCR tube, add 10-20 ng of the amplified library DNA from Step 10.35, adjust the total volume of the reaction to 10 µl, and incubate 30 min at 37°C. |
| | 3 | Add 18 µl (×1.8 volume) of AMPure XP beads, vortex mix, and wait 5 min at room temperature. |
| | 4 | Purify the DNA following Steps 11.16-11.22. |
| | 5 | Add 10 µl EB to the beads, vortex mix, and incubate 2 min at room temperature. |
| | 6 | Quick spin the sample tube, put on the magnet and collect the eluate in a new PCR tube. |
| | 7 | Analyze the size-shift of the library using 2 µl of the DNA with Agilent TapeStation High Sensitivity D1000 tape. Note: Place RE(-) and RE(+) samples for the same library side-by-side for easy comparison. |

##### Appendix: Reagents and consumables

- 1.5 ml Protein LoBind tube (Eppendorf, cat. 0030108116)

Note: For cell/tissue samples.

- 1.5 ml DNA LoBind tube (Eppendorf, cat. 0030108051)

Note: For DNA samples.

- 2.0 ml Protein LoBind tube (Eppendorf, cat. 0030108132)
- 50 ml tube (Thermo Fisher Scientific, cat. 14-432-22)
- 0.2 ml PCR tube (INA OPTICA, cat. 3247-00)
- 384-well PCR plate (Applied Biosystems, cat. 4309849)
- Optical Adhesive Film (Applied Biosystems, cat. 4311971)
- Liquid nitrogen
- Mortar and pestle (AS ONE, cat. 2-9037-02)
- SK mill (Tokken, cat. SK-200)
- Stainless-steel tube (Tokken, cat. TK-AM5-SUS)
- Stainless-steel bullet (Tokken, cat. SK-100-DLC10)
- Tube holder (Tokken, cat. SK-100-TL)
- Douncer (Sigma-Aldrich, cat. D8938)
- PBS minus (Wako Pure Chemical, cat. 314-90185)

Note: Make a 1X solution with H2O.

- 16% formaldehyde (Pierce, cat. 28906)
- Glycine (Wako Pure Chemical, cat. 077-00735)

Note: Make a 2.5 M solution with H2O.

- 1 M Tris-HCl (pH 8.0) (Wako Pure Chemical, cat. 314-90065)
- 5 M NaCl (Nacalai Tesque, cat. 31334-51)
- 0.5 M EDTA (Invitrogen, cat. 15575-038)
- 10% SDS (Invitrogen, cat. 15553-035)
- Triton X-100 (Sigma-Aldrich, cat. T8787)

Note: Make a 20% solution (w/v) with H2O.

- IGEPAL CA-630 (NP40) (Sigma-Aldrich, cat. 18896)
- Tween 20 (Sigma-Aldrich, cat. P9416)
- Proteinase inhibitor cocktail (Sigma-Aldrich, cat. P8340)
- Proteinase K solution (Nacalai Tesque, cat. 15679-64)
- RNase A (Takara, cat. U0505S)

Note: Make a 10 mg/ml solution with H2O.

- Phenol/Chloroform/Isoamyl alcohol (25:24:1) (Wako Pure Chemical, cat. 311-90151)
- Glycogen solution (Thermo Fisher Scientific, cat. R0561)
- 2-Propanol (Nacalai Tesque, cat. 03065-35)
- Ethanol (Junsei, cat. 17065-1230)

Note: Make 70% solution (for Steps 3 and 6.2) and 80% solution (for Steps 9, 11 and 12) with H2O.

- TE (pH 8.0) (Wako Pure Chemical, cat. 314-90021)
- EB (Qiagen, cat. 19086)
- Qubit dsDNA High Sensitivity Kit (Thermo Fisher Scientific, cat. Q32851)
- Agilent Bioanalyzer High Sensitivity DNA Kit (Agilent Technologies, cat. 5067-4626)
- Agilent TapeStation Genomic DNA ScreenTape (Agilent Technologies, cat. 5067-5365)
- Agilent TapeStation Genomic DNA Reagents (Agilent Technologies, cat. 5067-5366)
- Agilent TapeStation High Sensitivity D1000 ScreenTape (Agilent Technologies, cat. 5067-5584)
- Agilent TapeStation High Sensitivity D1000 Reagents (Agilent Technologies, cat. 5067-5585)
- NEBuffer 2 (New England Biolabs, cat. B7002S)
- NEBuffer DpnII (New England Biolabs, cat. B0543)
- DpnII (New England Biolabs, cat. R0543M)
- HindIII (New England Biolabs, cat. R3104M)
- ClaI (Takara, cat. 1034A)
- NheI (New England Biolabs, cat. R0131S)
- BSA solution (New England Biolabs, cat. B9000S)
- dNTP set (Thermo Fisher Scientific, cat. LS10297018)

Note: Make 1 mM solution (for Step 5) and 10 mM solution (for Step 8) with H2O.

- Biotin-14-dATP (Thermo Fisher Scientific, cat. 19524016)
- Biotin-14-dCTP (Thermo Fisher Scientific, cat. 19518018)

- Klenow DNA polymerase (New England Biolabs, cat. M0210L)
- T4 DNA ligase (New England Biolabs, cat. M0202M)
- T4 DNA polymerase (New England Biolabs, cat. M0203S)
- Covaris microTUBE (Covaris, cat. 520045)
- Agencourt AMPure XP beads (Beckman coulter, cat. A63880)
- Magnet stand for the microtube (Thermo Fisher Scientific, cat. 12321D)
- Magnet stand for a PCR tube (Nippon Genetics, cat. FG-SSMAG2)
- Streptavidin beads (Thermo Fisher Scientific, cat. 11205D)
- KAPA LTP Library Preparation Kit (KAPA Biosystems, cat. KK8230)
- KAPA HiFi HotStart Ready Mix (KAPA Biosystems, cat. KK2600)
- KAPA HiFi HotStart Real-time PCR Master Mix (KAPA Biosystems, cat. KK2701)
- Illumina TruSeq compatible adapter (PerkinElmer, cat. NOVA-514180)
- TPC mix (10 µM each)

Note: Mix two oligos; 5'-AATGATACGGCGACCACCGAG-3' and 5'-CAAGCAGAAGACGGCATACGAG-3'.

### Supplementary Protocol S2. Computational protocol to support multiple enzymes

#### *HiC-Pro*

**# Add restriction sites of Arima-specified enzymes to the `HiC-Pro/bin/utils/digest_genome.py` script**

```
RE_cutsite = {
    "mboi": ["^GATC"],
    "dpnii": ["^GATC"],
    "hinfi-1": ["G^AATC"],
    "hinfi-2": ["G^ATTC"],
    "hinfi-3": ["G^AGTC"],
    "hinfi-4": ["G^ACTC"],
    "bglii": ["A^GATCT"],
    "hindiii": ["A^AGCTT"]}
```

**# Run the script to make a restriction fragment file with multiple enzymes**

```
digest_genome.py -r dpnii hinfi-1 hinfi-2 hinfi-3 hinfi-4 -o HiC-Pro_arima.bed genome.fasta
```

**# Edit the HiC-pro `config-hicpro.txt` file for every combination of ligation sites**

```
LIGATION_SITE = ATCGATC,GAATGATC,GATTGATC,GAGTGATC,GACTGATC,GAATAATC,GAATATTC,GAATAGTC,
GAATACTC,GATTAATC,GATTATTC,GATTAGTC,GATTACTC,GAGTAATC,GAGTATTC,GAGTAGTC,GAGTACTC,GACTAAT
C,GACTATTC,GACTAGTC,GACTACTC,GATCAATC,GATCATTC,GATCAGTC,GATCACTC
```

**`LIGATION_SITE` should be written in one line*

##### *Juicer*

**# A python script for conversion of the restriction fragment file**

**hic-pro2juicer.py**

```
from signal import signal, SIGPIPE, SIG_DFL
signal(SIGPIPE,SIG_DFL)

import sys

output = ""
pre_chr = ""

# get HiC-Pro digestion bed file as the 1st parameter
args = sys.argv
path=args[1]

with open(path) as f:
    for s_line in f:

        # split line by tab
        line=s_line.split()

        # chromosome name
        chr = line[0]

        # cutting site
        site = int(line[2])
        site = str(site)

        if chr==pre_chr:
        # same chromosome
            output += " " + site
        else:
            # new chromosome
            if output != "":
                print (output)

            output = chr + " " + site

        # save chromosome name
        pre_chr=chr

# print the last chromosome
print (output)
```

**# Convert the restriction fragment file of HiC-Pro to the Juicer format**

```
python hic-pro2juicer.py HiC-Pro_arima.bed > Juicer_arima.txt
```
